# Single-cell spatial mapping reveals alteration of tissue microenvironment during early colorectal cancer

**DOI:** 10.1101/2024.11.20.622725

**Authors:** Tuhin K. Guha, Edward D. Esplin, Aaron M. Horning, Roxanne Chiu, Kristina Paul, Annika K. Weimer, Winston R. Becker, Rozelle Laquindanum, Meredith A. Mills, D. Glen Esplin, Jeanne Shen, Emma Monte, Shannon White, Thomas V. Karathanos, Daniel Cotter, Joanna Bi, Uri Ladabaum, Teri A. Longacre, Christina Curtis, James M. Ford, William J. Greenleaf, Michael P. Snyder

## Abstract

Familial adenomatous polyposis (FAP) is a rare, hereditary syndrome that raises the risk of developing colorectal cancer (CRC). This disease model is well suited for studying the early stages of malignant transformation. Our spatial CODEX experiments reveal that, in contrast to normal mucosa, FAP mucosa, pre-cancer polyps and colorectal cancers exhibit substantial alterations in the cell type composition and tissue microenvironment. These early alterations include: an increase in the population of cancer-associated fibroblasts (CAFs), and the inhibition of tumor infiltrated lymphocytes and cell-adhesion protein by CAFs, the transformation of memory T cells into regulatory T cells, nuclear translocation of beta-catenin from the cell membrane, a decrease in the M1:M2 macrophage ratio, a notable increase in angiogenesis events. Our studies define the early stem cell, stromal, and immune steps of colorectal cancer and may benefit early detection, and therapeutic intervention.

## Introduction

Oncogenesis involves a complex series of events, but the early events that occur during premalignant state formation and conversion to malignant tumors^1,2^ have not been well described. In particular, the cell types and the spatial organization of immune cells, cancer cells, stromal cells and cellular matrix components are poorly understood especially at the early stages of oncogenesis^3,4^. This information is important since cells with pre-cancer mutations are thought to be widespread and the cellular microenvironment is crucial for understanding conversion to oncogenesis and mechanisms of drug resistance.

Colorectal adenocarcinoma (CRC) is the third leading cause of cancer mortality in the U.S.^7^ and an ideal system to study early events in carcinogenesis. More than 90% of all CRC patients possess mutations that affect the Wnt signaling pathway, and more than 80% of these contain mutations in the Wnt-antagonist Adenomatous polyposis colitis (APC) gene which is thought to often be the initiating event for colorectal carcinogenesis. The APC protein acts as a scaffold in the beta (β)-catenin destruction complex thereby regulating the Wnt signaling pathway, which otherwise leads to intestinal hyperplasia^8–10^. Consistent with APC’s central role in the formation of polyps, individuals with germline inactivating mutations in APC exhibit the hereditary colorectal cancer syndrome known as Familial Adenomatous Polyposis (FAP) in which patients typically develop 100 or more polyps by early adulthood^11,12^. Thus, a single patient may present with multiple polyps of varying sizes and stages of development, all derived from a common germline lineage^13–15^. As such this is a useful model to study different stages of early carcinogenesis in the same original genetic background.

Previous studies have mapped cell types in sporadic polyps and those of FAP patients using single-cell RNA-sequencing (scRNA-seq) and assay for transposase-accessible chromatin with sequencing (ATAC-seq) and have detected cell types and transcriptional regulation programs within these polyps^16,17^. However, the differences with regard to the cellular spatial organization, composition of the functional neighborhoods, as well as the molecular changes within the tissue microenvironment (TME) of this disease continuum from healthy colon to invasive adenocarcinoma is not well characterized. In particular the location of the immune cells, cancer stem cells and stromal cells have not been mapped.

In order to understand the differences within TMEs during CRC, and as a part of the Human Tumor Atlas Network (HTAN)^18^, we have performed extensive spatial mapping of single cells from premalignant polyps, CRCs and normal mucosa (non-FAP healthy controls) using Co-Detection by indEXing (CODEX) single-cell multiplexed imaging platform^19,20^. Through spatial profiling we investigate the cell type composition and organization in normal mucosa and polyps of different stages including adenocarcinomas.

We observe an increase in stem-like cells, endothelial cells, cancer-associated fibroblasts (CAFs), regulatory T cells (Tregs), and M2 macrophages in early stage polyps when compared to healthy normal mucosa, suggestive of a suitable pre-cancer microenvironment for tumor cells to proliferate. In contrast, enterocytes, goblet cells, cycling Transit Amplifying cells (TA), and natural-killer cells (NK) decrease along the disease continuum. Compared to their counterparts in the FAP mucosa and normal mucosa, polyps and CRCs exhibit higher neo-angiogenesis events and strong nuclear expression of β-catenin. Within the glandular crypts of the FAP polyps, we find an increase in cancer stem cells (CSCs) as well as Tregs, tumor associated macrophages (TAMs), and vascular endothelial cells that potentially promote the survival and proliferation of CSCs. In the tumor “nest” of FAP adenocarcinoma samples, we find an immunosuppressive microenvironment where tumor cells are spatially separated from the immune cells. Additionally, we also observe that TAMs penetrate the tumorous region, promoting angiogenesis and tumor growth. We find that CAFs play a number of roles in supporting tumor growth and are typically found close to the inflammatory region within polyps and CRCs. Overall, our results provide insights into the complex series of events that occur during early stages of neoplastic formation which may ultimately be useful for therapeutic intervention and cancer prevention.

## Results

### CODEX generates single-cell spatial maps from FAP and non-FAP donor tissue samples

We generated CODEX single-cell spatial data from 52 samples (12 normal mucosa (from non-FAP patients), 16 FAP mucosa, 18 FAP polyps, 2 FAP colorectal adenocarcinomas and 4 sporadic colorectal cancers (CRCs)) collected across 12 FAP-affected individuals and 8 non-FAP individuals (Figure 1a). Of 20 donors, 12 were male and 8 were female, and the cohort consisted of 9 Hispanic, 10 European and 1 Black participant (Table S1). Age ranges were from 12 to 78 years. The samples were collected using colectomies. For comparison, normal mucosa samples were collected from deceased donors. The samples were collected from ascending, transverse, descending, and sigmoid regions of both FAP and non-FAP patient colons. (Table S2).

**Figure 1.**
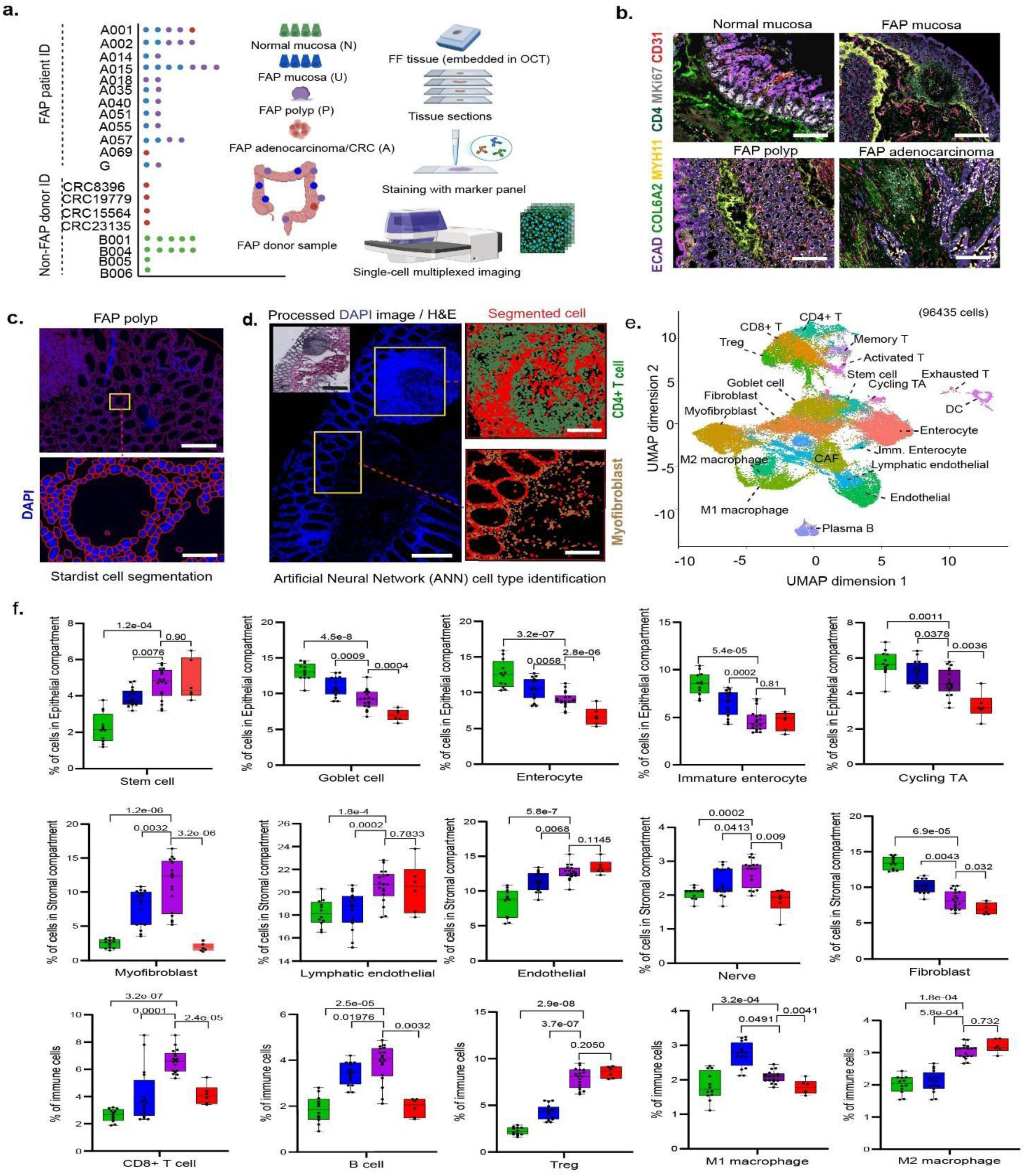
CODEX generates single-cell spatial maps of FAP and non-FAP donor samples and shows changes in cell type composition across malignant transformation. (a) FAP and non-FAP donor samples used in this study, Y-axis represents the donor IDs from 12 FAP patients and 8 non-FAP donors. The samples (colored dots) have been procured from different regions of the colon, flash-frozen and embedded in optimal cutting temperature (OCT) media for cryosectioning. The tissue sections were stained with a cocktail of barcoded-antibodies (marker panel) and highly multiplexed imaging was performed using Akoya’s PhenoCycler Fusion instrument. The samples have been color-coded throughout the text: Normal mucosa (green), FAP mucosa (blue), FAP polyp (magenta), FAP adenocarcinoma/sporadic CRC (red). (b) Representative images of normal mucosa, FAP mucosa, FAP polyp, and FAP adenocarcinoma stained with 40 antibodies [staining from 6 antibodies representing Epithelial layer (ECAD), fibroblast (COL6A2), myofibroblast (MYH11), CD4+ T cell, Proliferating cells (MKi67) and Endothelial/blood vessels (CD31) are shown here], scale bar: 100 µm. (c) Upper panel: Representative image of cell segmentation on DAPI-stained-nuclei (blue) from a glandular region of a FAP polyp section. The cell segmentation border is marked with red boundary, scale bar: 100 µm. Lower panel: Magnified view, scale bar: 20 µm. (d) Segmented cells (red) were further processed through Artificial Neural Network (ANN) pipeline for cell type identification; CD4+ T cells (green), myofibroblast (brown) are annotated and shown here as examples of such findings, scale bar: 200 µm, inset: 100 µm. (e) UMAP representation of all cell types detected. (f) Box plots depicting fraction of cells within epithelial compartment that are stem cells, goblet cells, enterocytes, immature enterocytes, cycling TA; within stromal compartment that are myofibroblasts, lymphatic endothelial, endothelial, cancer associated fibroblasts (CAFs), fibroblasts; and immune cell types such as CD8+ T cell, B cell, regulatory T cell (Treg), M1 macrophages and M2 macrophages, divided by disease state. Abundances of each cell type in normal mucosa, FAP mucosa and FAP adenocarcinoma/sporadic CRC are compared with their abundances in FAP polyp tissues with two-sided student’s *t* test and *P* values are listed in the plots. The boxplots are constructed from n = 12 normal mucosa, 16 FAP mucosa, 18 FAP polyp, and 6 FAP adenocarcinoma/sporadic CRC samples. Boxplots represent the median, 25^th^ percentile and 75^th^ percentile of the data; whiskers represent the highest and the lowest values within 1.5 times the interquartile range of the boxplot; and all points are plotted.

The tissues were analyzed using CODEX with up to 42 oligo-conjugated antibodies that were validated against different protein targets (Figure S1a, S1b). The majority of the cell type markers selected for CODEX imaging were based on single-nuclei RNA (snRNA) data that identified cell-specific gene expression information^17^. The antibodies were validated for specificity using protocols optimized for fresh frozen samples (see methods; Figure S2). The antibody marker panel (Table S3) was used to stain samples from normal mucosa, FAP mucosa, FAP polyp and adenocarcinoma/CRC patients (Figure 1b). The processed multiplexed images were segmented with StarDist cell segmentation pipeline (Figure 1c), (integrated within Qupath tissue analysis software) and both supervised and unsupervised Artificial Neural Network (ANN) were implemented (Figure 1d). Up to 25 cell types across the FAP disease continuum were identified using the oligo-conjugated antibody probes (Figure 1e).

### CODEX imaging shows changes in cell type composition across malignant transformation

The resultant dataset from the 52 spatial maps was used to quantify cell type composition in the normal mucosa and the FAP disease states. Overall, across the different disease states (FAP mucosa, FAP polyps, FAP adenocarcinoma, and CRCs) and including normal mucosa as controls, we observe an increase in stem-like cells, myofibroblasts, lymphatic endothelial cells, endothelial cells, cancer-associated fibroblasts (CAFs), B cells, CD8+ T cells, regulatory T cells (Tregs), exhausted T cells (present only in adenocarcinoma/sporadic CRC), and M2 macrophages. We observe a commitment decrease in enterocytes, immature enterocytes, goblet cells, transit-amplifying cells (TA), cycling TA cells, fibroblasts, and M1 macrophages (Figure 1f). Polyps collected from a 26-year-old and a 54-year-old FAP patient show similar immune cell type composition within the sub-mucosal intestinal lymphoid follicles (sm-ILF). For both cases, we observe dendritic cells (DC), CD4+ and CD8+ double positive T cells, PD1 marker expression within the germinal centers and vascular cells at the follicular periphery (Figure S1c). We also found that the percentage of stem-like cell populations within the crypts of normal mucosa decreases with increased age of the non-FAP donors (Pearson coefficient r = 0.9). However, even in FAP mucosa we observe both an increase in stem-like cells when compared with non FAP healthy mucosa, and a decreased correlation between the fraction of stem-like cells and the age of FAP patients (Pearson coefficient r = 0.2) (Figure S3). For example, a 62-year-old FAP patient had a similar fraction of stem-like cells to that of a 38-year-old FAP patient.

Increased immune function is strongly correlated with higher CD4+/CD8+ T cell ratio^24,25^ and a lower ratio is associated with a poor colorectal cancer prognosis^22,23^. We observe that the overall ratio of CD4+/ CD8+ T cells was significantly higher in normal mucosa (Figure S4a). We also observe increased numbers of CD4+ T cells localizing in the stromal compartment within the FAP mucosa when compared to FAP polyps. In contrast, the FAP polyps exhibit a relatively higher number of CD8+ T cells than CD4+ T cells and the CD8+ T cells tend to localize as intraepithelial lymphocytes (IELs) (Figure S4b).

### M2 macrophage is associated with FAP disease progression

Macrophages play a crucial role in maintenance of homeostasis, eradicating damaged cells and vascular tissue regeneration^30–33^. The two types of macrophages, M1 and M2, exist at variable levels depending on the microenvironment cues, and these cells tend to have different functions. For example, M1 macrophages can be tumor-resistant, whereas M2 macrophages promote tumor growth^34,35^ (Figure 2a). We observe a higher proportion of M1 macrophages within the normal and FAP mucosa. Conversely, the proportion of M2 macrophages were increased in FAP polyps and adenocarcinoma/CRC samples (Figure 2b). The M1: M2 macrophage ratio is an important determinant towards the prognosis of colorectal cancer and other intestinal bowel disorders (IBDs)^36,37^. We observe a higher M1: M2 macrophage ratio in normal mucosa and FAP mucosa and a lower ratio in FAP polyp and adenocarcinoma/sporadic CRC (Figure 2c). We also observe that M1 macrophages mostly colocalize with activated CD4+ T cells. Computation of the Pearson coefficient for colocalization between activated CD4+ T cells and M1 macrophages across all samples revealed significant correlation (R = 0.719) within normal and FAP mucosa when compared to that of FAP polyp and adenocarcinoma samples (P = 0.034; Figure 2d, e). This observation aligns with previous studies, where M1 macrophages have been shown to regulate T cell activation by providing the required cytokines and other costimulatory signals^38–40^.

**Figure 2.**
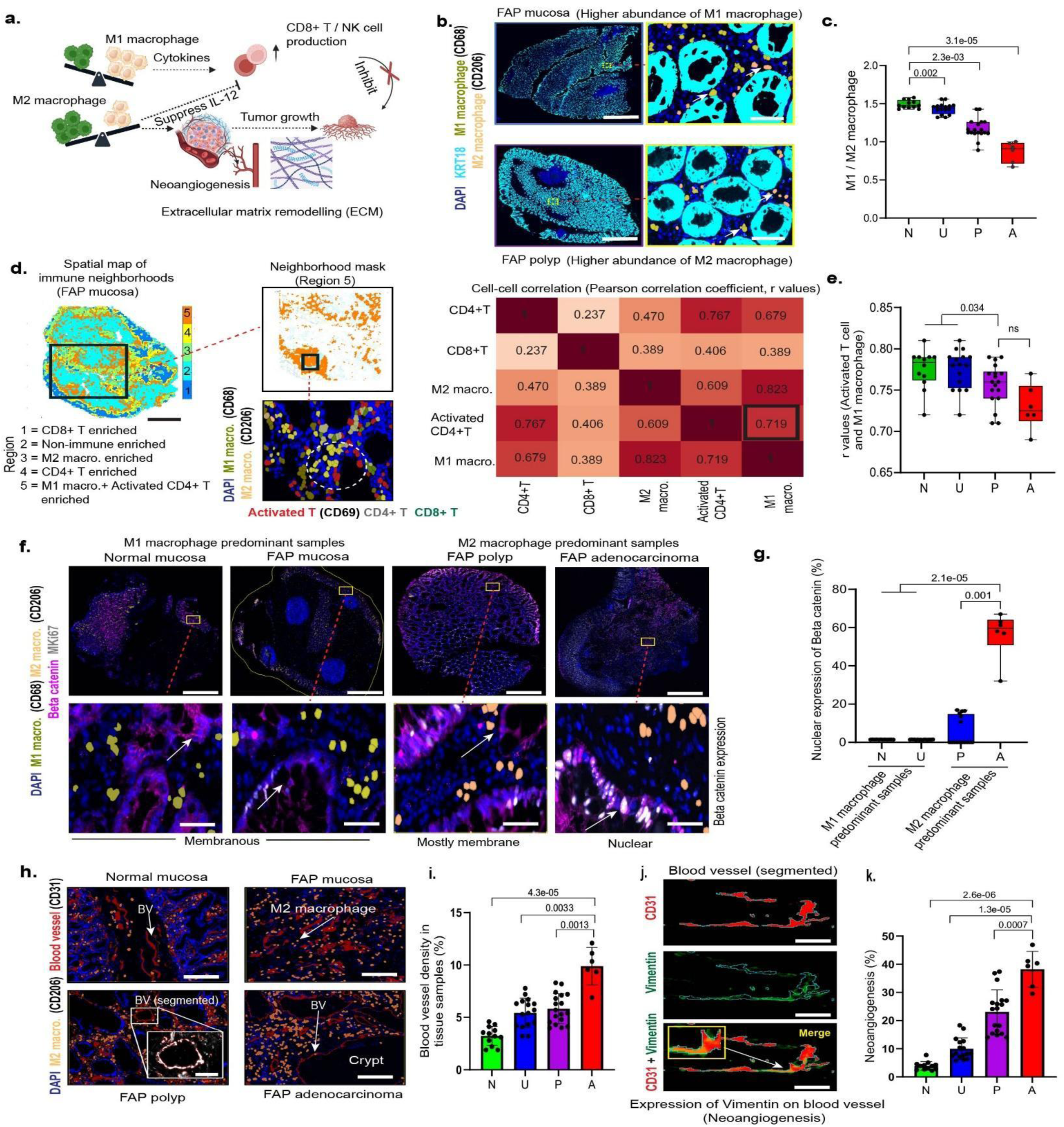
M2 macrophage is associated with FAP disease progression. (a) A schematic diagram showing the importance of M1 and M2 macrophage ratio within a tissue and its role within precancer microenvironment. M1 macrophages have antitumorigenic role while M2 macrophages have protumorigenic role (see text for details). (b) Representative images of FAP mucosa and FAP polyp showing the presence of M1 and M2 macrophages. Higher abundance of M1 macrophages is found in FAP mucosa, while relative abundance of M2 macrophages is higher in FAP polyp. The images have been magnified in the right panel for better visualization, scale bar: 500 µm, inset: 20 µm. (c) Boxplots depicting M1/M2 macrophage ratio in normal mucosa and across other FAP disease states. The ratio is compared with normal mucosa samples with two-sided student’s *t* test and *P* values are listed in the plots. The boxplots are constructed from n = 12 normal mucosa, 16 FAP mucosa, 18 FAP polyp, and 6 FAP adenocarcinoma/sporadic CRC samples. Boxplots represent the median,25^th^ percentile and 75^th^ percentile of the data; whiskers represent the highest and the lowest values within 1.5 times the interquartile range of the boxplot; and all points are plotted. (d) Representative image of a spatial map from FAP mucosa sample showing different regions/immune neighborhoods, scale bar: 200 µm. Region 5 with M1 macrophage along with activated CD4+ T cell enriched population has been shown as a neighborhood mask (orange) in the magnified image (top right). The annotated cell types comprising of M1 macrophage, M2 macrophage, activated CD4+ T cells, CD4+ T cells, and CD8+ T cells have been shown in the magnified image (bottom right). Heatmap showing cell-cell correlation between the above cell types, where M1 macrophage and activated CD4+ T cell show relatively higher cell-cell correlation (Pearson correlation coefficient, *r* = 0.719) compared to that of other cell types. (e) Boxplots showing Pearson correlation coefficient (*r*) values for colocalization of activated CD4+ T cell and M1 macrophage within normal mucosa and FAP disease states. Statistical student’s two-sided *t* test was performed and compared with FAP polyps. The boxplots are constructed from n = 12 normal mucosa, 16 FAP mucosa, 18 FAP polyp, and 6 FAP adenocarcinoma/sporadic CRC samples. Boxplots represent the median,25^th^ percentile and 75^th^ percentile of the data; whiskers represent the highest and the lowest values within 1.5 times the interquartile range of the boxplot; and all points are plotted. (f) Upper panel: Representative CODEX images showing membranous expression of beta-catenin in normal mucosa and FAP mucosa, where M1 macrophage is predominant than M2 macrophage. Representative CODEX images showing nuclear expression of beta-catenin in FAP polyp and FAP adenocarcinoma/sporadic CRC, where M2 macrophage is predominant than M1 macrophage, scale bar: 500 µm. Lower panel: Magnified images from each of the above representative images, scale bar: 20 µm. (g) Boxplots showing the nuclear expression of beta-catenin in M1 macrophage and M2 macrophage predominant samples. A two-sided student’s *t* test was performed on n = 12 normal mucosa, 16 FAP mucosa, 18 FAP polyp, and 6 FAP adenocarcinoma/sporadic CRC samples, and *P* values have been indicated. (h) Representative CODEX images of normal mucosa and FAP disease states, showing increase in M2 macrophages and blood vessel (BV) density across the disease continuum, scale bar: 100 µm. An example of a blood vessel (pixel segmented) has been magnified and shown within the inset, scale bar: 20 µm. (i) Bar plots showing percentage of blood vessel density in tissue samples. Statistical two-sided *t* test was performed and compared with adenocarcinoma samples. (j) Segmented blood vessel showing CD31 (red pixel mask, upper panel), vimentin (green pixel mask, middle panel) and merge (yellow pixel mask, lower panel) inferring the expression of vimentin on CD31 stained blood vessel (an example of neoangiogenesis event, see text for details), scale bar: 20 µm. (k) Bar plots showing the percentage of neoangiogenesis across the FAP disease states. A two-sided statistical student’s *t* test was performed and compared with FAP adenocarcinoma/sporadic CRC samples. N = Normal mucosa, U = FAP polyp, P = FAP polyp, A = Adenocarcinoma/CRC.

Beta-catenin is involved in both gene transcription and coordination of cell-cell adhesion^41,42^. Previous studies have shown that overexpression of β-catenin is associated with various cancers such as colorectal adenocarcinoma, lung cancer, malignant breast tumors, and ovarian cancer^43–45^. Moreover, alterations in the localization and nuclear expression levels of β-catenin (i.e. nuclear accumulation) have been associated with malignant transformation by activating several downstream target genes such as TCF, and LEF1^46–48^. Unlike M1 macrophages, M2 macrophage polarization promotes β-catenin expression in murine macrophage-like RAW264.7 cells and alveolar macrophages^49^. We observed membranous expression of β-catenin in normal and FAP mucosa, where the M1 macrophage population is higher than the M2 counterpart (Figure 2f). In contrast, we observed an increased nuclear expression of β-catenin in the FAP adenocarcinoma/sporadic CRC samples that display a higher population of M2 macrophages (Figure 2g).

Cancer cells, which require adequate supply of blood and nutrients, release angiogenic factors that encourage the growth of new blood vessels through a process called neo-angiogenesis. This process also involves vimentin, a type III filamentous protein, which helps in the migration, growth, and differentiation of endothelial cells lining the newly formed blood vessels^50,51^. Additionally M2 macrophages induced by IL-4, as well as neutrophils, express the highest level of pro-MMP9 which stimulates angiogenesis *in vitro*^52^. Consistent with these observations, we observe a significant increase in blood vessel density in the FAP affected tissues compared to normal mucosa (Figure 2h, i) and also observe vimentin protein expression along the endothelial lining of the newly formed blood vessels and microcapillaries (shown as branch formation), but not along mature blood vessels (Figure 2j,k).

### Regulatory T cells tend to localize near cancer stem cells needed for survival and proliferation

The normal colonic stem cell niche at the base of the glandular crypt is responsible for homeostasis and repair of the intestinal epithelial layer^53–55^. CRC stem cells (CSCs) have similar characteristics to those of normal colonic stem cells; however, these cells differentiate aberrantly, contributing to the supply of bulk tumor cells depending on several microenvironmental cues^56–58^. Consistent with this mechanism, we observe stem-like cells within the glandular crypts in both FAP mucosa and FAP polyp samples. Moreover, we detect a significant increase in the percentage of stem-like cells expressing a CSC marker, SOX2, in the FAP polyps compared to the FAP mucosa samples (Figure 3a).

**Figure 3.**
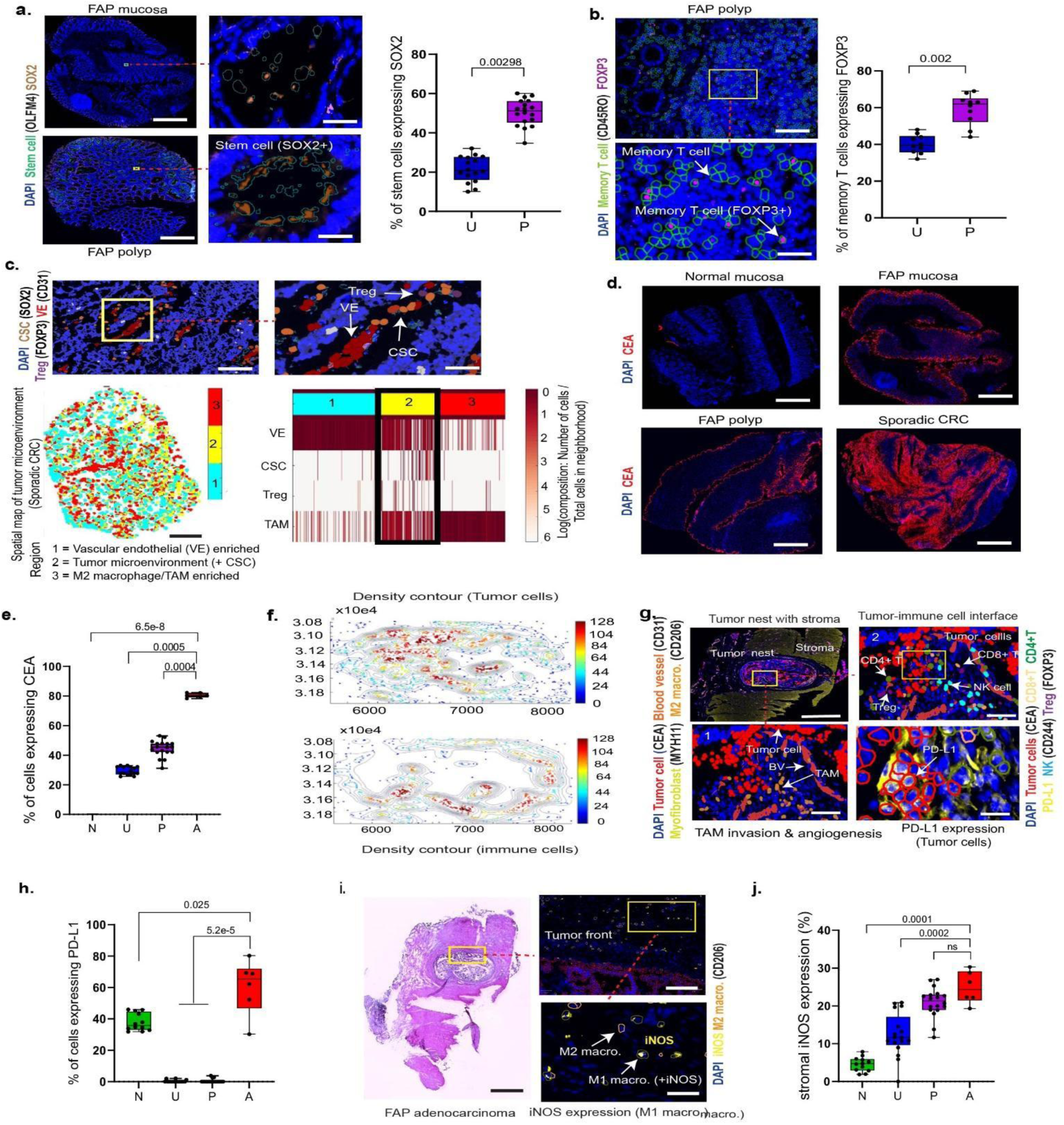
Regulatory T cells tend to be located near cancer stem cells during the initiation of precancer microenvironment. (a) Representative CODEX images of FAP mucosa and FAP polyp samples showing fractions of stem cells that express cancer stem cell marker, SOX2. FAP polyps have statistically higher percentage of stem cells expressing SOX2 than that of FAP mucosa. Box plots showing percentage of stem cells expressing SOX2 in FAP mucosa and FAP polyp. A two-tailed statistical *t* test was performed, scale bar: 200 µm, inset: 10 µm (b) Representative image of FAP polyp showing nuclear expression of FOXP3 (magenta color) within memory T cells (green boundary). Statistical t test was performed between FAP mucosa and FAP polyp samples, scale bar: 100 µm, inset: 50 µm (c) Representative CODEX image from a sporadic CRC showing the Treg and vascular endothelial (VE) located near the cancer stem cell (CSC), scale bar: 100 µm, inset: 50 µm. Spatial map of CRC microenvironment and heatmap showing region 1, 2 and 3, where region 2 comprise of VE, CSC, Treg, and tumor assisted macrophages (TAMs). (d) CODEX images showing increased expression of carcinoembryonic antigen (CEA) during the FAP disease progression, scale bar: 500 µm. (e) Box plot depicting percentage of cells expressing CEA marker in normal mucosa and FAP disease states. Statistical t test was performed and compared with FAP adenocarcinoma/CRC samples. (f) Density contour of tumor cells (upper panel) and immune cells (lower panel) within the tumor nest. The axes represent the cell coordinates and the intensity scale bar represents the density of the tumor cells (within the tumor nest) and immune cell types (along the tumor boundary) (g) CODEX image showing tumor microenvironment (tumor nest) from a FAP adenocarcinoma patient, specifically the tumor-immune interface, TAM invasion and angiogenesis. Tumor cells (segmented and shown with red boundary) expressing PD-L1 protein (yellow) on the surface, scale bar: 500 µm, inset: 10 µm. (h) Boxplots depicting the percentage of cells expressing PD-L1, statistical two-tailed t test was performed and compared with FAP adenocarcinoma (i) H&E staining of a FAP adenocarcinoma section, showing the tumor nest embedded within tumor stroma, scale bar: 500 µm. The magnified image (top panel) shows segmented cells and tumor front, scale bar: 100 µm, which was enlarged to show M1 macrophage expressing iNOS in the nucleus, scale bar: 20 µm. (j) Boxplots showing percentage of stromal iNOS expression across all sample types. A two-sided t test has been performed and compared with FAP adenocarcinoma/CRC samples. N = Normal mucosa, U = FAP mucosa, P = FAP polyp, A = FAP adenocarcinoma/sporadic CRC.

Regulatory T (Treg) cells expressing the Foxp3 transcription factor play a critical role in promoting the stemness of gastric cancer cells through the IL13/STAT3 pathway^59,60^ and also help in reducing overactive immune responses within the tumor microenvironment^61,62^. Within FAP polyps, we observe a significant increase in the percentage of memory T cells expressing Foxp3, indicating an overall increase in the Treg population (Figure 3b). In accordance with the previous findings that Tregs provide cytokines and interleukins (i.e. microenvironmental cues) for CSC survival and proliferation^63,64^, we also observe similar localization of Tregs in the FAP polyps near the same region as the CSCs (Figure 3c).

### Immunosuppressive microenvironment is found within FAP adenocarcinoma tumor **‘nest’**

Proliferation of transformed CSC-like cells ultimately allows malignant cells to expand from the intestinal crypt region and invade other surrounding tissue regions. Using a tumor marker, carcinoembryonic antigen (CEA), we observe increasing expression of CEA marker across the FAP disease continuum (Figure 3d), which was later quantified across all of the samples (Figure 3e). We also imaged a tumor ‘nest’ embedded within the stromal compartment of one FAP adenocarcinoma patient sample. Previous studies have shown that cells within the tumor ‘nest’ secrete angiogenic factors for tumor angiogenesis, which is needed for proliferation^65,66^. We detect tumor associated macrophages (TAMs) located at the tumor invasive front, consistent with observations that TAMs secrete various cytokines which contribute to the tumor angiogenesis process^67–69^. Moreover, our imaging data shows that the tumor ‘nest’ from the FAP adenocarcinoma sample creates an immunosuppressive microenvironment by excluding most of the immune cells, such as CD8+ T cells, CD4+ T cells, B cells, and NK cells. However, these immune cells have been detected outside the periphery of the tumor ‘nest’. Moreover, density contour maps (a particular kind of 2-D density map, which represents the density of data points (representing cells) using contour lines)) of both CEA+ tumor cells within the ‘nest’ (Figure 3f, upper panel) and immune cells located outside the ‘nest’ have been generated (Figure 3f, lower panel). We also observe expression of Programmed Death Ligand 1 (PD-L1) on the tumor cells (Figure 3g) and the expression of this protein has been found only in normal mucosa and FAP adenocarcinoma/sporadic CRC samples, and not in FAP mucosa and pre-cancer polyps (Figure 3h). Studies have found that M1 macrophages express inducible nitric oxide synthase (iNOS) presumably as part of the macrophage inflammatory response^70–72^. In our CODEX imaging data, we also observe M1 macrophages (surrounding the tumor ‘nest’) expressing iNOS protein within the nuclei (Figure 3i) and there is an increase in the stromal iNOS expression across the FAP disease continuum (Figure 3j).

### Cancer associated fibroblasts (CAFs) play an essential role in FAP disease progression

The stromal compartment, comprising basement membrane, fibroblasts, extracellular matrix, immune cells, and vasculature, maintains both normal epithelial tissues and their malignant counterparts. It has been shown that changes within the stromal microenvironment at the tumor invasion front lead to the transdifferentiation of fibroblasts to cancer associated fibroblasts (CAFs). Several tumor-derived cytokines such as transforming growth factor-β (TGF-β), epidermal growth factors are thought to be responsible for this transformation. Activated CAF within the polyp and tumor microenvironment also has a positive effect on the formation of new blood vessels (angiogenesis) and M2 macrophage polarization, and may assist in extracellular matrix remodeling^75^ (Figure 4a). We detect inflammatory fibroblasts (involved in the inflammatory response in general; stained with Podoplanin (PDPN)), and CAFs (specifically associated with cancer development and progression; stained with Versican (VCAN)) within the precancer and tumor stroma, across the FAP disease (Figure 4b). In particular, the expression of PDPN and VCAN is higher in adenocarcinoma/sporadic CRCs (Figure 4c). We further inquired if CAFs are present near the inflammatory site and potentially responsible for the inflammation in the FAP affected colorectal tissue samples. Both FAP polyps and tumor samples were stained with VCAN (CAF marker; green), and PDPN (inflammatory marker; red), which were imaged (Figure 4d). To quantify the overlap (red pixels + green pixels = yellow pixels), a surface plot of the marker intensity was generated for each of these markers followed by computing the Vansteelsen’s cross correlation coefficient (CCF)^80^, which measures how the Pearson coefficient changes after shifting the red image voxel over green image voxel by different distances. We find that the overlap CCF between CAF marker (VCAN) and Inflammatory fibroblast marker (PDPN) increases while the overlap CCF between CAF marker (VCAN) and normal fibroblast marker (COL6A2) decreases in FAP adenocarcinoma/CRC samples (Figure 4e). This indicates that CAFs are present in close proximity to the inflammatory region within the FAP affected tissues adenocarcinomas/sporadic CRCs.

**Figure 4.**
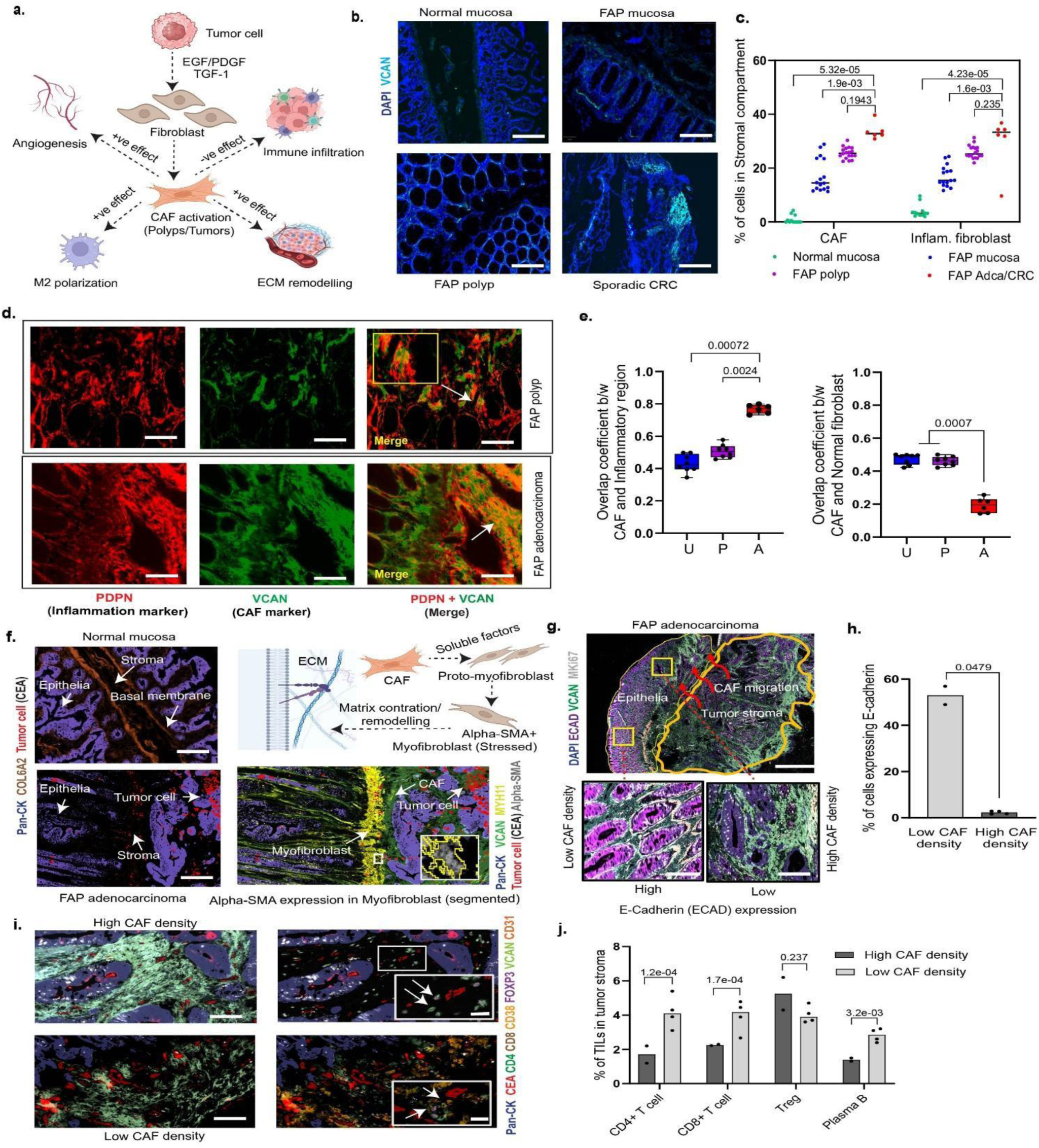
Cancer associated fibroblasts (CAFs) play an essential role in FAP disease progression. (a) Schematic representation showing the role of CAFs in tumor progression by positively regulating angiogenesis, M2 macrophage polarization, extracellular matrix remodeling and negatively regulating immune cell infiltration towards the tumor area. (b) Representative images of CODEX images from normal mucosa, FAP mucosa, FAP polyp and FAP adenocarcinoma showing increase in the expression of CAF (VCAN marker) across the disease continuum, scale bar: 100 µm. (c) Dot plots showing the percentage of CAF and inflammatory fibroblast within the stromal compartment from all sample types. Two sided statistical t tests were performed for each cell type and P values have been indicated. (d) Representative marker staining showing the overlap of PDPN (inflammation marker) and VCAN (CAF marker) within FAP polyp and FAP adenocarcinoma, scale bar: 100 µm. (e) Box plots showing overlap coefficient between CAF and inflammatory region (left panel) and overlap coefficient between CAF and normal fibroblast within FAP mucosa, FAP polyp and FAP adenocarcinoma/sporadic CRC. Two-sided t tests have been performed and compared with the tumor samples and P values have been indicated. (f) Representative CODEX images showing presence of basal membrane in normal mucosa and absence of the same in FAP adenocarcinoma, scale bar: 100 µm. Upper right panel: Schematic diagram showing how CAF influences extra cellular matrix remodeling by modifying myofibroblasts through alpha-SMA expression within myofibroblasts. Lower right panel: Representative CODEX image from a FAP adenocarcinoma sample showing the alpha-SMA expression (white) within myofibroblasts (segmented with yellow boundary). (g) Representative CODEX image of FAP adenocarcinoma showing the migratory route (red curved arrows) of CAF expanding from tumor stroma towards epithelial glandular region (mucosa), scale bar: 500 µm. Two different regions are magnified to show higher expression of E-cadherin in glandular cells near the region with low CAF density and lower expression of E-cadherin near the region with high CAF density, scale bar: 50 µm. (h) Bar plot showing percentage of cells with FAP adenocarcinoma and sporadic CRC samples, expressing E-cadherin at low CAF density and high CAF density. Two tumor samples were found with low CAF density and four tumor samples with high CAF density. A two-tailed statistical t test has been performed between the samples and the p value has been indicated. (i) CODEX images show the high and low CAF density from FAP adenocarcinoma and sporadic CRC sample respectively, scale bar: 50 µm. Tumor sample with high CAF density shows lesser infiltration of lymphocytes (or tumor infiltrated lymphocytes/TILs) while low CAF density shows higher TILs in the tumor stroma, scale bar: 10 µm. (j) Bar plots showing the percentage of TILs such as CD4+ T cell, CD8+ T cells, Tregs and plasma B cells within high CAF and low CAF density from the FAP adenocarcinoma and sporadic CRC samples. Statistical t tests were performed for each immune cell type and P values have been indicated.

CODEX imaging also revealed how stromal CAFs invade the epithelial layer in FAP tumor samples leading to further spreading of tumors. We observe a continuous lining of the basal membrane in normal mucosa while FAP tumor samples have disrupted or minimal basal membrane layer (Figure 4f, left panel). Cancer associated fibroblasts (CAFs) secrete soluble factors which turn proto-myofibroblasts into myofibroblasts. These myofibroblasts under stressed conditions express alpha-smooth muscle actin (α-SMA) protein which results in extracellular matrix contraction/remodeling^81^. We observe similar expressions of α-SMA protein from the myofibroblasts (Figure 4f, right panel). Moreover, it has been observed from our imaging data that CAFs also affect the expression of E-cadherin (ECAD), which is a membrane protein that helps to maintain the epithelial barrier by joining neighboring cells together to form tissues^82^. We observe less E-cadherin expression in glandular epithelial cells along the CAF migratory path as compared to the glandular cells further away from the CAFs migratory route, which continue to express higher levels of this cell-cell adhesion protein (Figure 4g), which were quantified (Figure 4h).

Tumor infiltrating lymphocytes (TILs) are an integral part of the body’s immune system, identifying and eliminating abnormal cells, including cancer cells^83,84^. In our study, we observe a higher CAF population within a FAP adenocarcinoma sample. However, the corresponding stromal area within that same tumor sample with a higher CAF density has been detected with a lower number of TIL population (Figure 4i). In contrast, we see an inverse correlation between the number of CAFs and the presence of TILs because sporadic CRC samples with lower CAF density show a higher number of TIL population (Figure 4i). Overall, the percentage of TILs comprising CD4+ T cell, CD8+ T cell, and plasma B cells decreased with increase in CAF population. Interestingly, high CAF density seems to have no effect on the migration or presence of the regulatory T cells (Treg) within both high and low CAF density tumor stroma (Figure 4j).

### Multicellular neighborhood analyses from FAP mucosa and adjacent FAP polyp show differences in neighborhood composition

To understand how intercellular interactions, cellular densities, and multicellular structures change across the disease continuum, we performed cell neighborhood analyses^88^ (Figure 5a). Heatmap reflecting the enrichment of cell types preferentially co-localizing revealed 16 significant multicellular structures, which were broadly classified as epithelial, stromal and immune-based neighborhoods. Six neighborhoods comprising glandular epithelial, CEA+ cell enriched epithelial, neutrophil enriched epithelial, secretory epithelial, mature epithelial and CD8+ intraepithelial regions were classified as epithelial neighborhoods. Seven neighborhoods comprised of endothelial cells, stromal mucin pool, innate immune cell population, MYH11+ myofibroblast enriched, CAF enriched, microstructures (blood vessels), and macrostructures (lymphatic endothelial cells) were classified as stromal neighborhoods. The rest of the neighborhoods, composed of inner follicle, outer follicle and innate immune cell hub, were classified as immune neighborhoods (Figure 5b).

**Figure 5.**
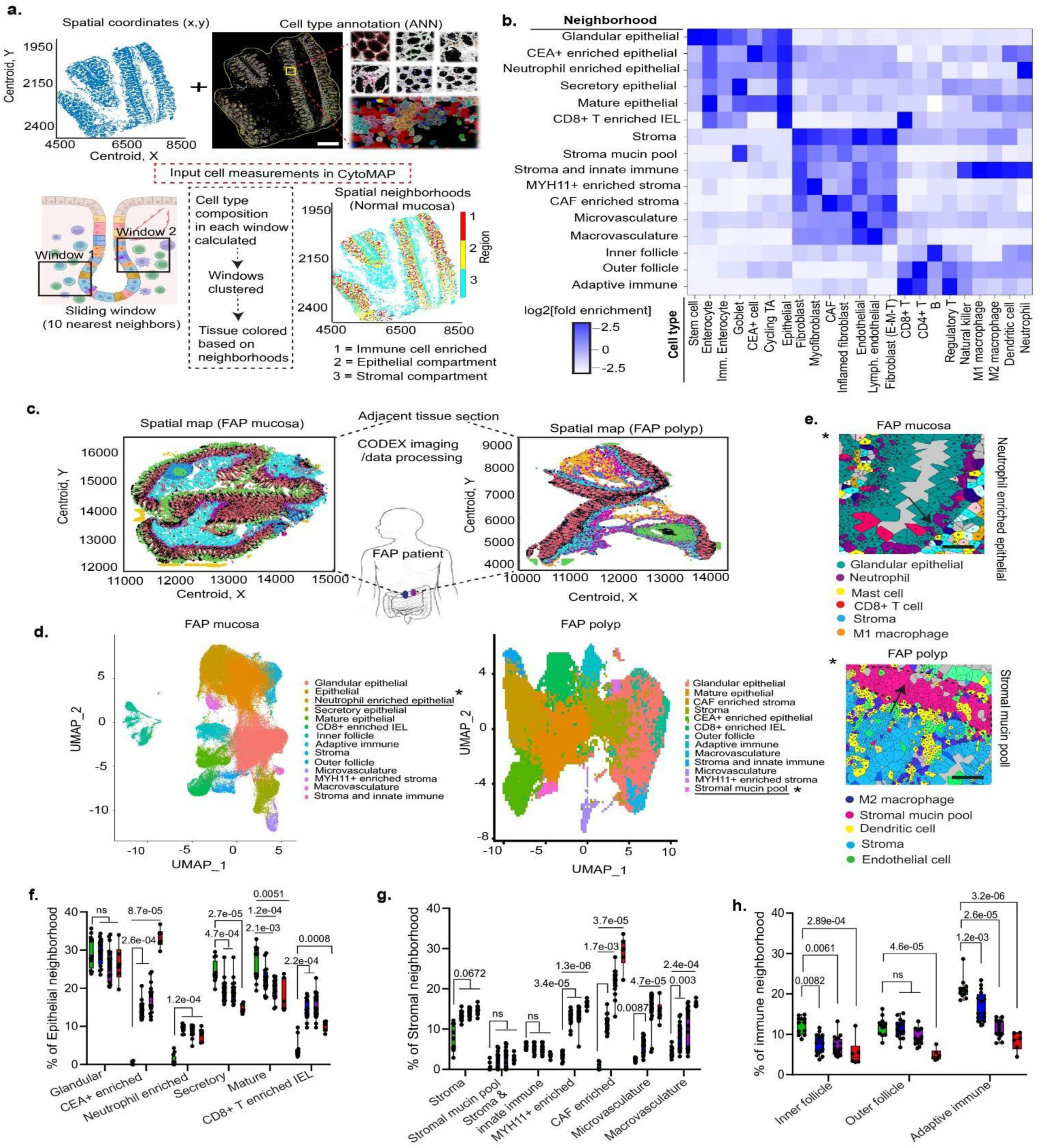
Cell neighborhood analyses reveal alteration of tissue microenvironment during precancer events. (a) Schematic representation of cell neighborhood analysis, where spatial coordinates and cell type annotation (cell measurements) are used as input for CytoMAP pipeline. Neighborhood analysis was done by taking a window across cell types and vectorizing the number of cell types in each window, clustering windows, and assigning clusters as cellular neighborhoods. (b) Sixteen unique intestinal multicellular neighborhoods were defined by enriched cell types as compared to the overall percentage of cell types in the samples. (c) Representative spatial maps of adjacent FAP mucosa and FAP polyp samples from the same patient (d) UMAP projection of cell neighborhood clusters from FAP mucosa and FAP polyp samples have been generated. Neutrophil-enriched from FAP mucosa sample and stromal mucin pool from FAP polyp have been indicated with asterisks. (e) Upper panel: Voronoi representation of glandular epithelial cells (green) showing the presence of neutrophils (magenta) constituting the neutrophil-enriched epithelial neighborhood. Lower panel: Voronoi representation of stromal mucin pool (pink) within FAP polyp sample. (f) Epithelial neighborhood percentage as a percentage of epithelial neighborhoods from multicellular analysis performed on each sample separately. (g) Stromal neighborhood percentages as a percentage of immune neighborhoods from multicellular analysis performed on each individual sample separately. (h) Immune neighborhood percentage as a percentage of immune neighborhoods from multicellular analysis performed on each individual sample separately. Statistical t tests were performed for each neighborhood and compared with respect to the normal mucosa samples and the respective P values have been indicated.

To understand the spatial co-association of cells in the pre-cancer microenvironment and its transformation during FAP disease progression, we carried out multicellular neighborhood analyses on adjacent FAP mucosa and FAP polyp sample pairs. Representative spatial maps of a pair of adjacent FAP mucosa and FAP polyps from the same patient are shown (Figure 5c). The neighborhoods from these tissue sections were clustered (Figure 5d) revealing similar neighborhood compositions during the disease progression from FAP mucosa to FAP polyp, the FAP mucosa. However, we observe a new, neutrophil-enriched epithelial neighborhood, secretory epithelial neighborhood and an inner follicle composed of mainly B cells. Our observation of neutrophil-enriched epithelial neighborhoods in FAP mucosa (Figure 5e) has been reported in other intestinal bowel disorders (IBDs) and has been associated with severity of disease symptoms and disruption of critical barrier function and neutrophil-epithelial interaction^89^.

### Tissue microenvironment changes across FAP disease continuum

Multicellular neighborhood analyses revealed differences in the structural composition across the FAP disease continuum as well as in the composition of these neighborhoods. In the epithelial compartment, we observed no change in glandular epithelial neighborhood across the FAP diseased states when compared with the normal mucosa. However, the percentages of CEA+ cell-enriched, neutrophil-enriched neighborhoods increased, while the secretory and mature epithelial neighborhoods decreased within the epithelial compartment, across the FAP malignancy continuum. Previously, it has been reported that intraepithelial CD8+ lymphocytes (IELs) are located in the spaces between intestinal epithelial cells and are the first to interact with pathogens that cross the epithelial barrier^91^. IELs are essential for immune homeostasis regulation and epithelium integrity maintenance^92,93^. Congruent to the previous findings^94,95^, we observe increased CD8+ T cell intraepithelial neighborhoods in the precancer states compared to normal counterparts. Interestingly, this IEL neighborhood decreased in FAP adenocarcinoma/sporadic CRC samples (Figure 5f).

More stroma is produced during malignancy, actively promoting tumor development, invasion, and metastasis^96,97^. In our spatial data, we observe significant increases in stromal neighborhood, MYH11+ enriched myofibroblast neighborhood, CAF enriched neighborhood along with microvasculature and macrovasculature neighborhoods within FAP precancer and tumor microenvironment compared to that of the normal mucosa tissue samples (Figure 5g).

At least two different multicellular neighborhoods exhibited changes in CD4+ T cell prevalence and localization. The membership of CD4+ T cells, B cells, and dendritic cells defined two distinct follicle-based structures. CD4+ T cells were more abundant in the first of these structures, which are found in the follicle’s outer regions, while B cells were more abundant in the follicle’s inner regions. A fully developed lymphoid follicle was required for the inner follicle neighborhood, also known as the germinal center (Figure S1c). We observe an overall decrease in inner follicle structures across the FAP diseased states, while the outer follicle structures decreased only in the adenocarcinoma samples (Figure 5h). An adaptive immune neighborhood consisting of CD8+ T cells also decreased across the disease continuum. This may be due to a progressive loss of adaptive immune infiltrate and by the establishment of a progressively immune “cold” microenvironment^98,99^.

## Discussion

Premalignant events lead to abnormal changes responsible for triggering cancer precursors that favor the initiation of a tumor^100,101^. Therefore, studying the precancer environment is crucial for earlier diagnosis, novel interventions, and risk stratification thereby improving tumor risk management and leading to potential prevention strategies. Previous research on colorectal cancer biology mainly focused on advanced-stage tumors^102–105^ and has largely disregarded pre-cancer events. Even though snRNA and ATAC sequencing help understand differences between individual cells, cellular heterogeneity and complex biological systems, both these techniques dissociate the cells, thereby losing important spatial information and context. Our CODEX multiplexed imaging fills the gap by spatially mapping single cells across the FAP disease continuum, thereby providing information about how cells organize and interact across the pre-cancer microenvironment and inform key changes during the disease progression. A summary of the main findings has been presented (Figure 6).

**Figure 6.**
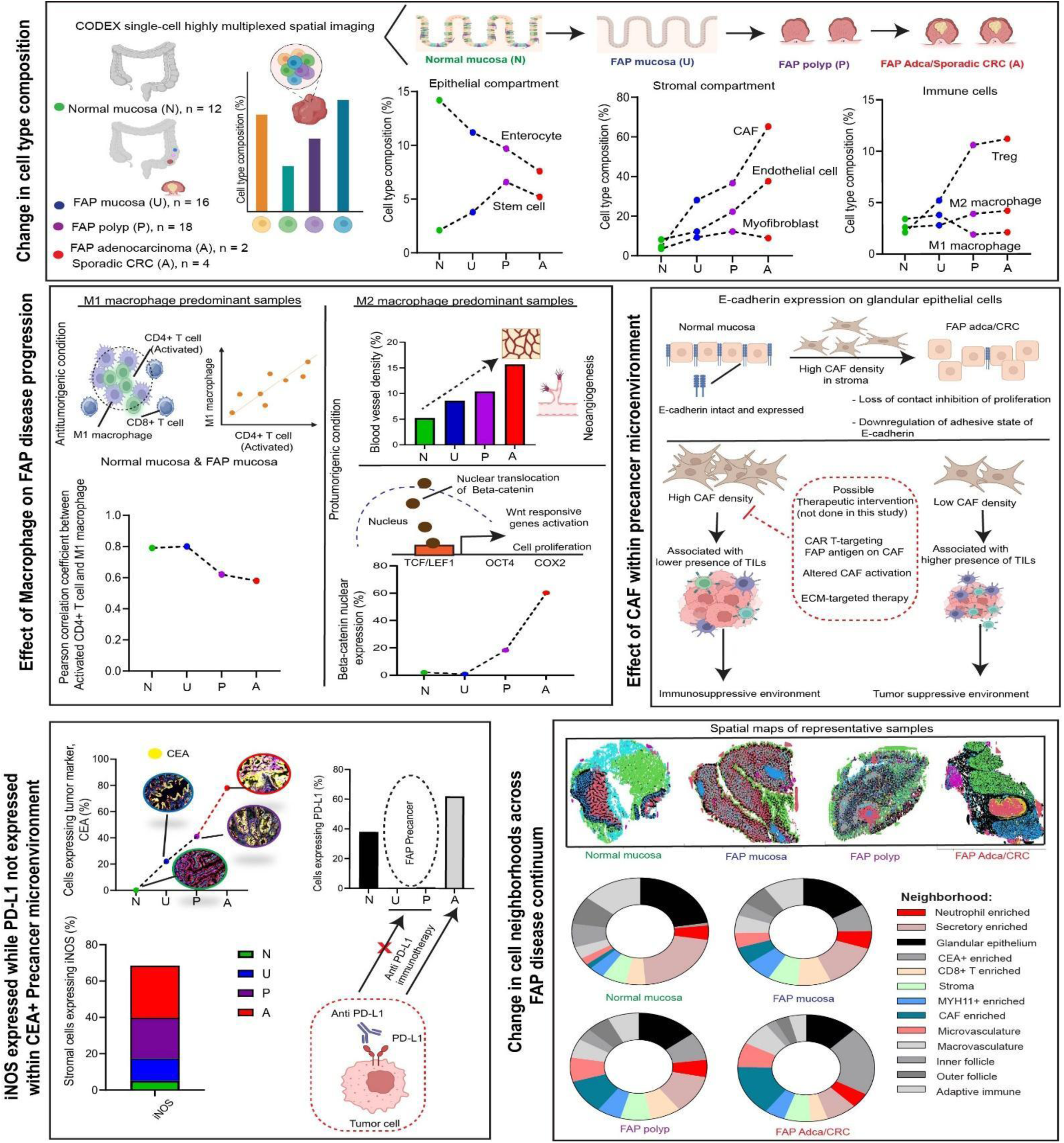
Summary of the main findings. Discoveries were classified into five main groups: change in cell type composition, effect of macrophage on FAP disease progression, effect of CAF within precancer microenvironment, expression of iNOS and absence of PD-L1 proteins within precancer polyps and change in cell neighborhoods across FAP disease continuum.

Analysis of cell type composition reveals an increase in stem-like cells, CAFs, Tregs, M2 macrophages and decrease in NK cells, M1 macrophages, goblet cells across the disease continuum. A pre-cancer microenvironment that encourages the growth of cancer cells is probably favored by the observed changes in cell type composition across the disease continuum. For example, the decrease in goblet cells in intestinal crypts and increase in CAFs, Tregs and exhausted T cells in the stroma overall favor an immunosuppressive environment within premalignant polyps and adenocarcinoma. Identifying the similar cell types within normal and FAP mucosa shows different cell type composition, which are unfavorable for tumor growth. The CD4+/CD8+ ratio is an important indicator of the overall strength of the immune system within a patient’s body. Higher CD4+ T cells over CD8+ T cells contribute to a relatively stronger immune system, while the reverse indicates an impaired immune response, potentially making it challenging for the body to effectively fight off tumor progression^24,25^. Within the immune cell population, we observe a decrease in CD4+/CD8+ T cell ratio during the FAP disease progression. The CD8+ T cells positioned at the interface of the intestinal lumen of the colon and external environment not only help in balancing immune tolerance but also protect the fragile intestinal barrier from invasion^26^. It could also be possible that patients with higher CD4+/CD8+ ratio (as seen in FAP mucosa compared to FAP polyp and FAP adenocarcinoma/CRC) have a better clinical course, with significantly higher 5-year survival^24,25^.

We also find that the increase in stem-like cell population within pre-cancer polyps has no correlation to donor’s age. This is expected because self-renewal capabilities of stem-like cells divide to create more stem cells in advanced polyps, which is essential for maintaining stemness and tumor formation. However, signals from other cell types in the vicinity also determine whether stem cells will self-renew, differentiate or remain quiescent. In our study, we find Tregs localize near stem-like cells within FAP polyps and CRCs, which play a critical role in promoting the stemness, possibly by providing cytokines and interleukins for the CSC survival and proliferation. Vascular endothelial cells and TAMs have also been observed near the CSCs, which further creates a favorable environment for stem cell proliferation. Recent studies involving single-cell RNA sequencing of gastric cancer and adjacent tissues found that Tregs were recruited to gastric cancer tissues and have further demonstrated that Tregs promote the stemness of gastric cancer cells through the IL13/STAT3 pathway^59,60^. As a result, we find an increase in the fraction of stem-like cells expressing cancer marker, SOX2, within polyps compared to FAP mucosa. Therefore, it would be interesting to see if blocking the interaction between Tregs and CSCs within the pre-cancer microenvironment could be a potential approach in the treatment of CRC. Within the immune microenvironment, both resident and recruited macrophages are key modulators during tumor progression which have several immunoregulatory functions. Similar to CD4+:CD8+ T cell ratio, M1:M2 macrophage ratio also decreases along with the FAP disease continuum. We find more M2 polarization occurring in advanced polyps than normal and FAP mucosa. This is expected as M1 is anti-tumorigenic and M2 is pro-tumorigenic^34^. Moreover, we observe a higher M2 polarized macrophage population that tends to favor nuclear expression of β-catenin, which is normally found in the epithelial cell membrane, where it helps with cell-to-cell adhesion. In the nucleus, β-catenin is associated with transcription factors from the TCF/LEF family and drives transcription of Wnt/β-catenin target genes leading to malignancy^41^. Previously it has been shown that mRNA/protein levels of β-catenin, Axin2, and c-Myc were significantly increased in M2 macrophages compared with that in M0 or M1 macrophages^97^. Hence, interfering with the macrophage polarization during the pre-cancer stage could be a turnaround step preventing nuclear localization of β-catenin, thereby curbing transcription of malignancy-related genes.

Solid tumors require sufficient blood supply to grow beyond a few millimeters in size. It has been shown that nearby normal cells are also stimulated by tumors to produce angiogenesis signaling molecules^65,66^. We find an increase in vasculature and microcapillaries within FAP polyps and tumors compared to normal and FAP mucosa. Most studies have reported angiogenesis once the invasive adenocarcinoma has been established^98–101^. However, we find that even in premalignant stages, such as FAP mucosa and FAP polyps, epithelial cells have increased proliferation and therefore would be expected to require increased blood supply. Therefore, targeting blood vessels in the premalignant stage with angiogenesis inhibitors such as, Axitinib (Inlyta®), Bevacizumab (Avastin®),Cabozantinib (Cometriq®) that specifically recognize and bind to vascular endothelial growth factor (VEGF) could be an effective route to interrupt pre-cancerous polyp formation and progression.

Among the other fibroblasts that we have detected within the stromal compartment of pre-cancer polyps and tumors, CAFs have been a central component of the tumor microenvironment, as they not only interfere with cell-cell adhesion but also interact with cancer cells via secreted molecules, influence cancer cells via extracellular matrix (ECM) remodeling and immune cell infiltration^102^. We have shown that CAFs are present near the inflammatory regions and inhibit the expression of E-cadherin protein, thereby interrupting the cell-cell adhesion feature. Moreover, CAFs have been shown to influence immune infiltration. Our imaging data shows that higher CAF populations in the stromal regions are only able to limit the entry of CD4+ T cells, CD8+ T cells, and plasma B cells into the tumor microenvironment, which results in the creation of an immunosuppressive environment. Usually, high stroma is associated with poor disease prognosis^96^. Therefore, both CAF and high stromal content might have restricted the immune cell infiltration. Despite the fact that CAFs have long been thought to be a key factor in the development of cancer and thus make an appealing therapeutic target, most clinical trials that attempt to target CAFs have failed^103,104^. Though the current research focuses on finding new CAF subsets that promote tumors and ways to specifically target them, it’s important to note that finding tumor-suppressive CAF populations and how they maintain their homeostatic balance will also be valuable as future stroma-targeted therapies. Consequently, therapeutic approaches with the potential to improve patient survival include normalizing or re-engineering the tumor stroma into a quiescent state or even tumor-suppressive phenotypes would be beneficial.

Since it can be used to comprehend the cellular makeup of the tissue and find prospects for extra in-depth investigation, cellular neighborhood analysis is a good place to start for a variety of downstream analyses. Once the dataset’s cell types or states have been annotated, we may determine if these annotations are spatially enriched. Therefore, we can find clusters that are neighbors in the tissue of interest by computing a neighborhood enrichment. To put it simply, it’s an enrichment score based on the spatial closeness of clusters: data from one cluster will appear to be enriched and have a high score if they are frequently adjacent to observations from another cluster. However, the score will be low if they are infrequent neighbors due to their distance from one another^88,105^. Using cell neighborhood analyses, we have found 16 different cellular neighborhoods across the FAP disease states. Within the epithelial compartment of FAP mucosa, we observed an increase in the neutrophil-enriched neighborhood. Understanding the molecular basis and functional implications of neutrophil-epithelial interactions has long been of interest because migration of neutrophils across mucosal epithelia is linked to disease symptoms and disruption of critical barrier function in conditions like inflammatory bowel disease^106–108^. We observe an increase in intraepithelial CD8+ lymphocytes (IELs) neighborhood within the pre-cancer microenvironment. Because they provide immune surveillance against pathogens, CD8+ T cells within the intestinal epithelium (IEL) are essential for preserving the integrity of the gut mucosa. However, their dysregulation can result in inflammatory bowel disease (IBD) and other intestinal disorders, making their clinical implications important for understanding and treating these conditions^109,110^. In IBD, particularly Crohn’s disease, an increased number of activated CD8+ IELs can be observed within inflamed intestinal mucosa, contributing to tissue damage and inflammation^111^. Adenocarcinoma/sporadic CRC samples have shown a decrease in the CD8+ IEL neighborhood. This could be attributed to the fact that tumor cells create an immunosuppressive microenvironment and also limit migration or infiltration of CD8+ T cells. The population of CD8+ IELs is not uniform, and based on their phenotypic traits and tissue location, various subsets may have unique roles^112^. New treatments that modify the activity of CD8+ IELs to reduce inflammation or improve immune protection may result from a better understanding of the distinct subsets and signaling pathways involved in CD8+ IEL function^113^.

Overall, our single-cell spatial imaging data demonstrates the changes in cell type composition throughout initiation of polyp formation from mucosa and the malignant transition of polyps into CRC, offering insights into the organization and interactions of cells inside the pre-cancerous milieu and driving important modifications during the course of the disease. We also discussed a few potential interventions based on these insights that could be deployed within the pre-cancer microenvironment, in order to curb tumor initiation and progression. This research is expected to have a significant impact on understanding the FAP pre-cancer microenvironment, as well as early stages of sporadic CRCs and other gastrointestinal malignancies, which will facilitate the implementation of early treatment and potential preventive interventions.

## Method details

### Tissue collection and processing

The Stanford University Institutional Review Board (IRB) and the Washington University Institutional Review Board (normal mucosa from deceased organ donors) have both approved this study, which conforms with all applicable ethical regulations. For every donor participant, the next-of-kin provided written informed consent. All of the FAP patient tissues used in this study were procured from colons surgically resected via total colectomy. The patient colons were taken directly from the surgical suite to the Stanford Hospital pathology gross room, where they were quickly rinsed and bisected with surgical scissors longitudinally on a room temperature cutting board typically used in a pathology gross room. Polyp-adjacent normal mucosa (absent any visible polyps) and polyp tissues were carefully dissected from the colon, measured, stored in cryovial tubes, and flash frozen by placing the samples in cryotubes directly into liquid nitrogen. Polyps that had a maximum diameter in any direction that was larger than 10 mm were considered “large”, between 10 mm and 5 mm were “medium”, and less than 5 mm were considered “small”. All polyps greater than 10mm in diameter were frozen sectioned, H&E stained, histopathologically examined by project pathologists to assess percentage of normal vs dysplastic tissue and presence of dysplasia by grade (low grade dysplasia vs high grade dysplasia). Polyps less than 10mm in size were not uniformly subjected to this histopathologic analysis, due to the low likelihood of high-grade dysplasia in this size of polyp and the need to preserve the very limited amount of tissue available for downstream multi-omic analysis. To record the exact location in the colon from which the different tissue samples were obtained for patients A001, A002, A014, A015, we marked the location of each tissue sample with a numbered thumb-tack and took photographs of the bisected and pinned-open colonic lumen. Using a ceramic mortar and pestle, we pulverized the flash frozen samples in liquid nitrogen to create a fine powder or small chunks as input to each multi-omic assay. For a subset of tissue samples, flash frozen polyps and colorectal mucosa were embedded in frozen OCT media and serial sectioned for 2D structural analyses. Once flash frozen or embedded in frozen OCT media, tissue was preserved at −80 degrees C until utilized for cryosectioning and CODEX imaging experiments.

### CODEX antibody conjugation and antibody panel creation

CODEX multiplexed imaging was carried out using the staining and imaging protocol outlined in the manufacturer’s user manual. Detailed step-by-step protocol for antibody custom conjugation has been shared in protocols.io https://www.protocols.io/edit/protocol-for-custom-conjugation-of-oligo-barcodes-g3k4bykyx Antibody panels were designed to include targets that distinguish intestinal epithelial and stromal cell subtypes, as well as innate and adaptive immune system cells and cancer cells. Detailed panel information is provided in Table S3. Each antibody was conjugated to a unique oligonucleotide barcode, and the tissues were stained with the antibody-oligonucleotide conjugates following the manufacturer’s protocol. We verified that the staining patterns in positive control tonsil or intestine tissues matched those previously established for immunohistochemistry. Prior to being evaluated collectively in a single CODEX multicycle, antibody-oligonucleotide conjugates were first evaluated with immunofluorescence assays, where the signal-to-noise ratio, antibody dilution and other technical aspects such as automated microfluidics and microscope were assessed.

### Preparing tissue samples for CODEX imaging

Imaging data were gathered from twenty human donors, each of whom represented a dataset. Each dataset contains tissue samples from pre-cancer polyps and adenocarcinoma/CRC collected from different sections of the colon (sigmoid, descending, transverse and ascending). The tissues were individually frozen in Optimal Cutting Temperature (OCT) molds and then sectioned at a width of 5-8μm using a cryostat, assembled on Leica Superfrost adhesive slides and histological staining was performed and/or stored at −80 °C, until needed. Histological staining was performed following the protocol, which has been shared in protocols.io https://www.protocols.io/edit/hematoxylin-and-eosin-staining-h-amp-e-protocol-fo-g3ktbykwp The tissue slides were processed according to the manufacturer’s protocol. The detailed protocol can be found in protocols.io https://www.protocols.io/edit/protocol-for-staining-fresh-frozen-tissue-sections-g3k2bykyf Briefly, the slides were obtained from the freezer on the day of the CODEX experiment and placed on 1-2 cm Drierite beads for 2 mins, followed by 10 mins incubation in acetone. The acetone was dried by placing the slides facing up in a humid chamber for 2 mins. The tissue sections were incubated in a 5 ml Hydration buffer for 2 mins (twice) and fixed using 1.6% Paraformaldehyde (PFA) for 10 mins, after which the fixative was thoroughly removed by rinsing in the same Hydration buffer. The slides were incubated in a 5 ml staining buffer for 20 mins. In the meantime, a stock solution of CODEX blocking buffer was prepared by mixing 362 µl of Staining buffer and 9.5 µl each of the company’s proprietary buffers namely N, G, J, S blockers (per 2 samples). The volume of antibody per sample slide was calculated and subtracted from the CODEX blocking buffer. An antibody cocktail solution was prepared by pipetting each antibody into the CODEX blocking buffer. The sample slides were placed in the humidity chamber and the antibody cocktail staining solution (200 µl per tissue sample) was dispensed on the slide and incubated for 3 hours at room temperature.

Subsequently, unbound antibodies were removed by rinsing the slides in a 5 ml Staining buffer for 2 mins (twice). The samples were placed in a post-staining fixative solution (1 ml of 16% PFA + 9 ml of Storage buffer) for 10 mins and washed thoroughly with 1x Phosphate buffer saline (PBS) to remove the fixative. Subsequently, the tissue sections were fixed with ice cold methanol for 5 mins at 4 °C and washed with 1x PBS. The samples were incubated in a final fixative reagent (20 µl of CODEX fixative reagent + 1 ml of 1x PBS) for 20 mins, washed thoroughly with 1x PBS before placing in the storage buffer (up to 2 weeks at 4 °C). In the meantime, a reporter plate for interrogating the corresponding barcoded-antibodies was prepared by adding nuclease-free water, 10x CODEX buffer, assay reagent, and nuclear stain. Two forty microliters of the stock solution were dispensed into individual wells followed by 5 µl of the corresponding reporter dyes. The reporter solutions were mixed and pipetted into individual wells of a 96-well plate, covered with a foil seal, and stored at 4 °C for imaging. The procedure to prepare reporter plates has been shared in protocols.io https://www.protocols.io/edit/preparing-the-phenocycler-reporter-plate-g3k5byky7

### CODEX multiplexed imaging

A CODEX microfluidic instrument also known as PhenoCycler Fusion (Akoya Biosciences Inc., CA, USA) integrated with an inverted fluorescence microscope through a custom stage insert was used to automate CODEX buffer exchange and image acquisition. The tissue slides were placed on the stage insert and imaged with multiple cycles of CODEX imaging. The first and the last cycles were basically blank cycles and also used for image registration and image alignment. Tissue sections were imaged in a 5x7 tiled acquisition at 377 nm/pixel resolution and 9 z-planes per tile. The tile overlap was preset at 30%.

### CODEX data processing and visualization

The image data and experiment.json files were transferred to CODEX Analysis Manager (CAM) for further processing of the raw images, which were then subjected to background subtraction, shading correction, along with deconvolution to remove out-of-focus light using a Microvolution software available from http://www.microvolution.com/. After drift-compensation and stitching, the best focal plane of vertical image stacks collected at each acquisition were chosen for cell segmentation using a gradient-tracing watershed algorithm (Akoya built-in software) on the nuclear staining (radius set to 6 pixels). The processed images were visualized and analyzed using CODEX Multiplex Analysis Viewer (MAV) installed as an extension within ImageJ software (https://imagej.nih.gov/ij/). MAV enables visualization, annotation, and analysis of cell populations from CODEX imaging data. For each cell in the imaged tissue, MAV generates spatial coordinates and measures the integrated signal intensity for each antibody. Occasionally, while using manual gating, some cell segmentation noise that could affect cell type identification have been revealed but using Leiden-based unsupervised clustering, followed by over-clustering and manually overlaying the resultant cell type clusters to the image resulted in much accurate identification of cell types. Both CODEX MAV and Qupath software can be downloaded from https://help.codex.bio/codex/mav/installation/download-and-install and https://qupath.github.io/. Alternatively, Qupath platform was also used for image data visualization and performing cell measurements followed by histo-cytometric multidimensional analysis pipeline (CytoMAP) to analyze the processed images (https://gitlab.com/gernerlab/cytomap). Once the paired.csv files were saved in Qupath, the annotated cell for every dataset was imported into CytoMAP. All of these include the spatial locations, gate definitions, and cell statistics for every cell object. This spatial analysis technique extracts and quantifies information about preferential cell-cell associations, global tissue structure, and cellular spatial positioning using a range of statistical techniques.

### Cell type analysis

The identification of cell types was carried out using previously described methods^20,105^. Briefly, DAPI+ cells were gated to select nucleated cells, and then protein markers used for clustering were normalized using z-normalization. Then the data were overclustered with X-shift clustering function within CODEX MAV program or Leiden-based clustering embedded within the Seurat package (version 5.1.0.). Seurat performs uniform manifold approximation and projection (UMAP), computing cell clusters and scales the data to observe variations in antigen expression on cells. The above steps were performed based on an in-depth analysis of marker intensity associated with cell types and its localization within the tissue and referred to the published images for that specific antibody. Based on the location within the image and the average cluster protein expression, a cell type was assigned to each cluster.

### Cell neighborhood analysis

Using raster scanned neighborhood analysis, CytoMAP determines the local cell composition inside a spherical (3D data) or circular (2D data) area/volume in the tissue. This function will treat the data as effectively 2D and use a cylindrical neighborhood window if the z dimensionality of the data is non-zero but the z thickness is less than the neighborhood radius. This function determines the total number of cells in each neighborhood as well as the maximum fluorescence intensity (MFI) of each channel. With half of the user-defined radius separating the neighborhood centers, the neighborhoods are uniformly spaced out in a grid pattern throughout the tissue. Then, additional analysis (e.g., local cellular densities, cell-cell associations) can be performed using the neighborhood data.

### Classify neighborhoods Into regions

The cell neighborhoods have been further grouped into regions. Multiple types of neighborhood information are used to define tissue regions using CytoMAP’s *region* function. This comprises the composition (number of cells divided by the total number of cells in each neighborhood), the raw number of cells per neighborhood, and the standardized number of cells of each phenotype in each neighborhood (number of cells minus mean number of cells, divided by the standard deviation of the number of cells in each neighborhood across the dataset). The number of regions and the neighborhood clustering were automatically determined by default using the Davies-Bouldin function. Once the network parameters were determined, the network was trained on the neighborhood data using MATLAB’s *train* function. This assigns a cluster number to each neighborhood. At the beginning of the selforgmap algorithm, NR “neurons” are scattered throughout the data. Then, in order to match the data’s landscape, it iteratively shifts the neurons’ positions closer to the data. In this case, the position is topological within the cell composition data rather than spatial. Next, each neighborhood’s closest neuron is located, leading to the clustering of the neighborhoods. Using CytoMAP’s Edit Region Colors function, the individual regions’ arbitrary color designations were altered for visual aids. With CytoMAP’s Heatmap Visualization feature, the color-coded neighborhoods’ composition was plotted. By creating a new figure in CytoMAP, plotting the neighborhood positions, and choosing the regions for the ‘c’ axis to color-code the neighborhoods according to region type, the spatial distribution of the regions was made visible.

### Cell-Cell Correlation Analysis

Certain cell populations preferentially associate with one another, or conversely avoid one another, based on the local cell density within individual neighborhoods, which can be used to correlate the location of different cell types. The number of cell or object types within the scanned neighborhoods was calculated using the Pearson correlation coefficient by the correlation, or corr function, within CytoMAP. This information is then graphed on a heatmap plot. This correlation analysis can be carried out on a number of samples, covering entire tissues or just specific areas of the tissues. This is significant because different tissue compartments may have different associations between cells.

### Resource availability Lead contact

Further information and requests for resources and reagents should be directed to and will be fulfilled by the lead contact, Prof. Michael P. Snyder

### Materials availability

Correspondence and requests for materials should be addressed to Prof. Michael P. Snyder. All biological materials can be obtained from the corresponding authors following reasonable requests.

### Data and code availability

Raw and processed CODEX image data generated in this study have been deposited to Synapse (www.synapse.org/) and are indexed on the Human Tumor Atlas Network (HTAN) Data Portal. The list of study files can be retrieved by accessing the HTAN Data Portal (https://data.humantumoratlas.org/explore) and filtering for Atlas: “HTAN Stanford” AND Assay: “CODEX”. Codes for processing the single-cell CODEX imaging data using Akoya’s pipeline such as CODEX MAV and Qupath can be downloaded from https://help.codex.bio/codex/mav/installation/download-and-install and https://qupath.github.io/ respectively. CytoMAP scripts for analyzing the processed images and neighborhood analysis can be found at https://gitlab.com/gernerlab/cytomap. Seurat pipeline for analyzing the single-cell imaging data can be found at https://cran.r-project.org/web/packages/Seurat/index.html.

## Acknowledgements

This work is supported by NIH grants no. U2C CA233311 (M.P.S., J.M.F., C.C., W.J.G.), no. 1U54HG010426 (M.P.S., W.J.G.), no. RM1-HG007735, no. UM1-HG009442, no. UM1-HG009436, no. R01-HG00990901 and no. U19-AI057266 (to W.J.G.). Y. Lin of Washington University in St. Louis provided the B001, B004, B005 and B006 HuBMAP healthy colon tissue samples (for comparison with the FAP affected tissues). BioRender was used for generating Figs.1a, 2a, 4a, 4f.

## Author contributions

T.K.G., M.P.S., J.M.F., and W.J.G. conceptualized the study. T.K.G. generated the histology slides, performed CODEX multiplexed imaging, analyzed the data and generated the figures. R.L., M.M., A.M.H., U.L., E.D.E., A.K.W., E.M., K.P., R.C., S.W., J.B. were responsible for sample collection. A.M.H., A.K.W. and E.M. were involved in project coordination. W.R.B. assisted in preparing the original CODEX panel. T.L. and J.S. performed histopathologic assessment of the samples. D.C. and T.K. handled data management and submission to repositories. T.K.G., M.P.S. and W.J.G wrote the original draft of the manuscript. All authors reviewed and edited the manuscript. C.C., E.D.E., J.M.F., M.P.S. and W.J.G. were responsible for funding acquisition. M.P.S. and J.M.F. supervised the project.

## Declaration of generative AI and AI-assisted technologies

During the preparation of the manuscript and associated figures (and supplemental figures), we did not use any generative AI and AI-assisted technologies.

## Supplemental information

**Figure S1:**
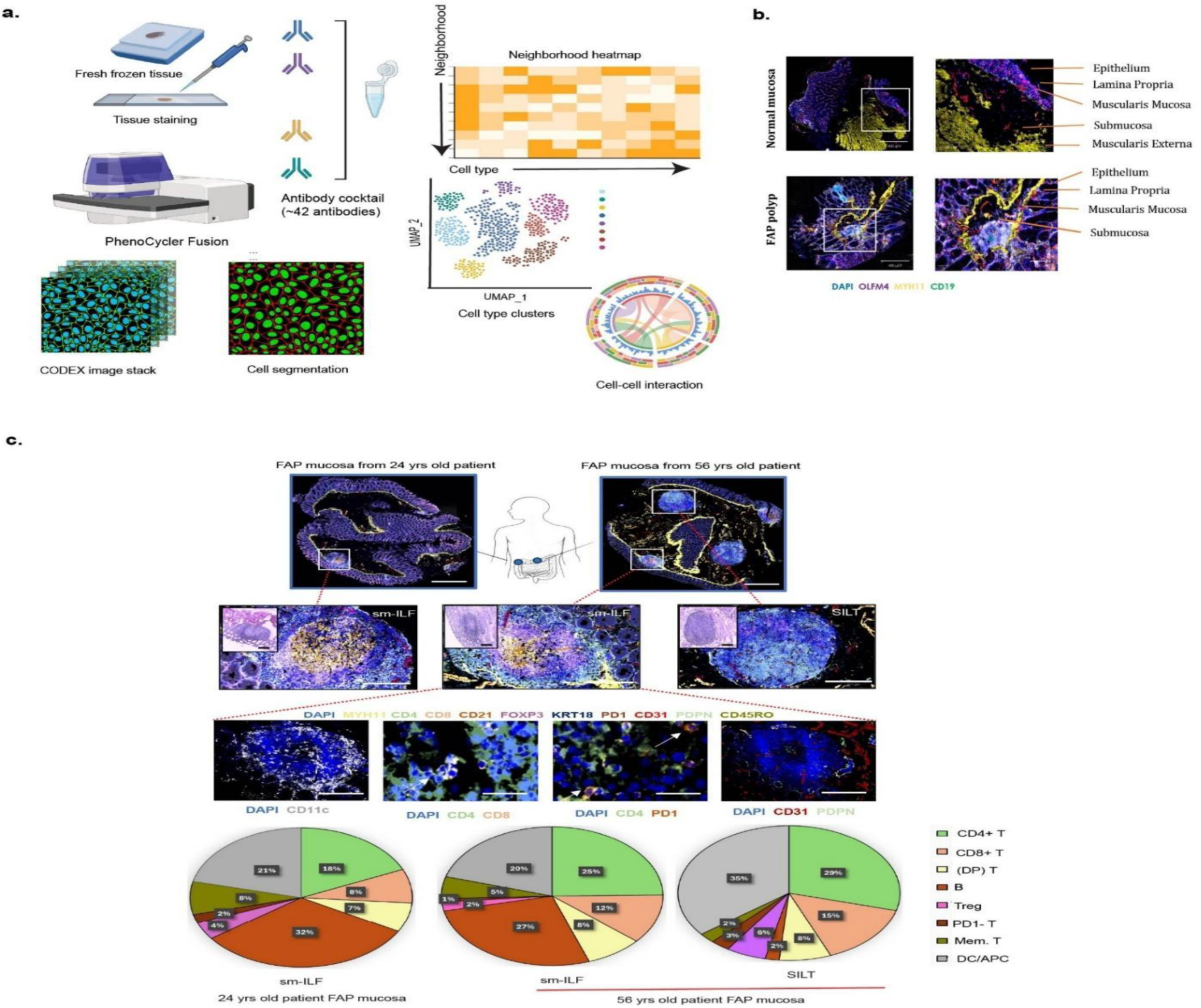
Detecting overall colon tissue morphology and lymphoid follicles using standard CODEX workflow. (a) Schematic representation of CODEX workflow (see text for details). (b) Representative CODEX images showing overall tissue morphology from normal mucosa (upper panel) and FAP mucosa (lower panel), scale bar: 200 µm (c) CODEX images of FAP mucosa samples from 24 year old and 56 year old FAP patients, showing the presence of submucosal intestinal lymphoid follicles (sm-ILF) and solitary intestinal lymphoid tissue (SILT), scale bar: 200 µm. CODEX detects the presence of dendritic cells (DC), CD4+ T cells, CD8+ T cells, Double-positive (DP)-T cells and microcapillaries within the sm-ILF. Lymphatic endothelial cells are present in the periphery, scale bar: 50 µm. Pie charts representing the percentages of various immune cell types within these lymphoid structures have been provided.

**Figure S2:**
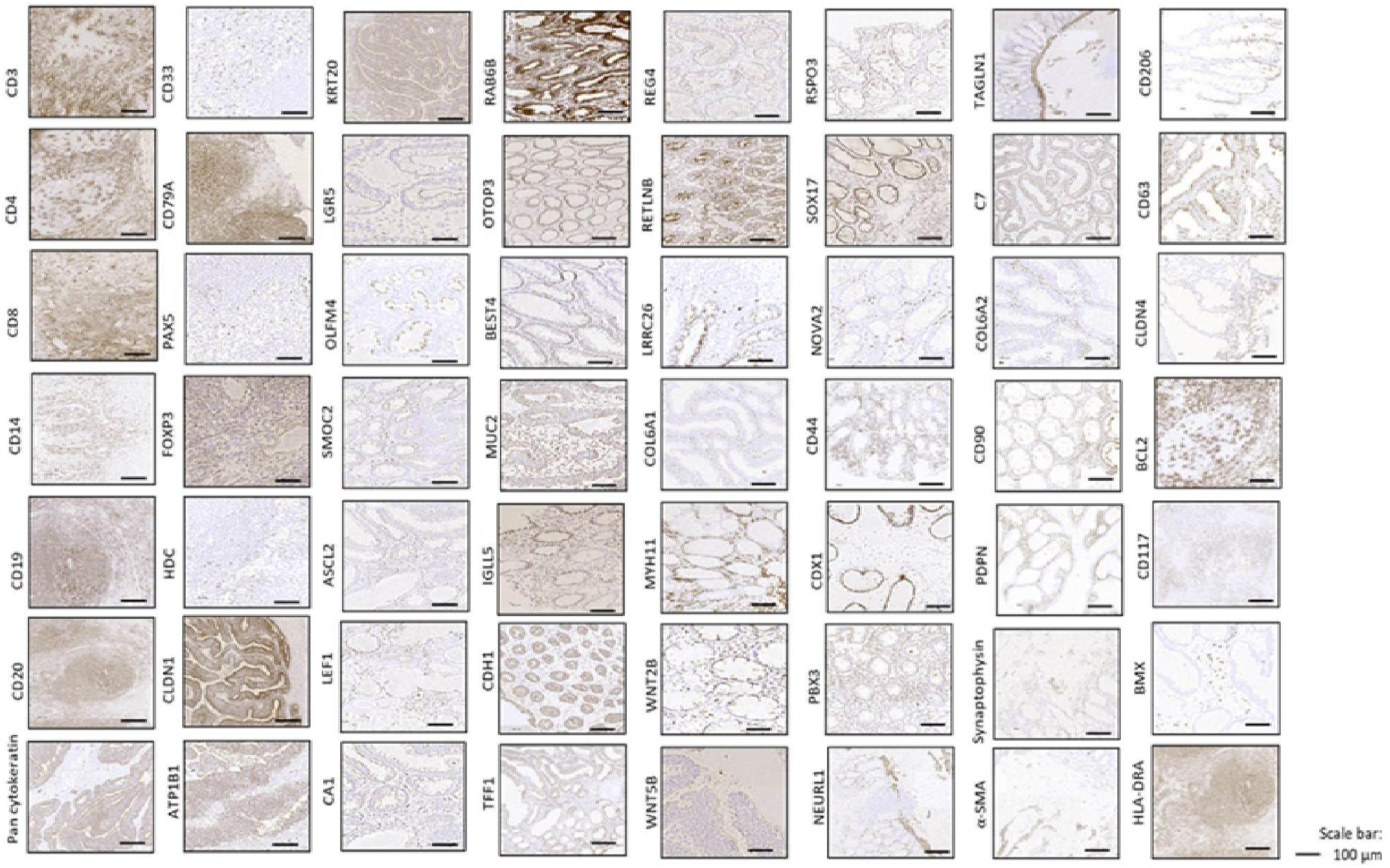
Antibody validation. Individual antibody validation performed on positive control tissues and colon tissues before including in the CODEX marker panel for performing CODEX experiments.

**Figure S3:**
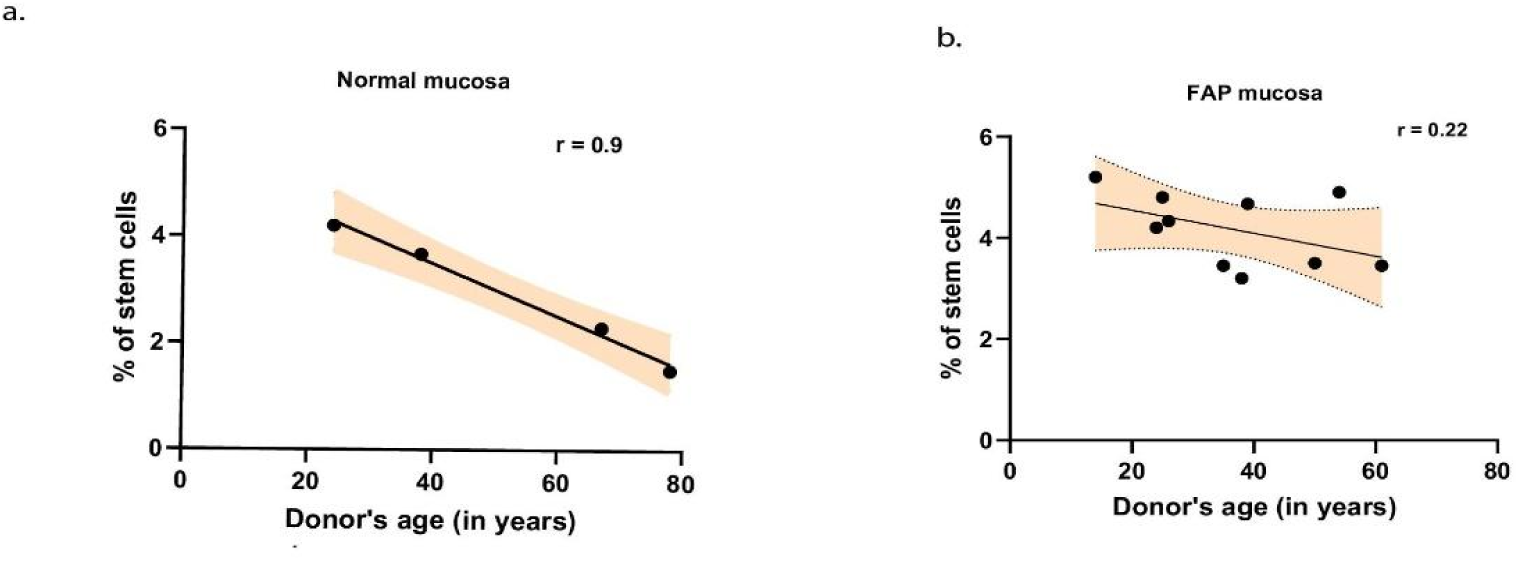
Correlation of stem cells found in polyp samples and donors’ age. Correlation of intestinal stem cell population with donor’s age from (**a**) normal mucosa: Strong correlation has been observed (r = 0.9) indicating stem cells decrease with increase in age within normal mucosa samples (**b**) no such correlation has been observed (r = 0.22) between age and stem cell population derived from FAP mucosa samples, indicating stem cells proliferate irrespective of the age of the patient.

**Figure S4:**
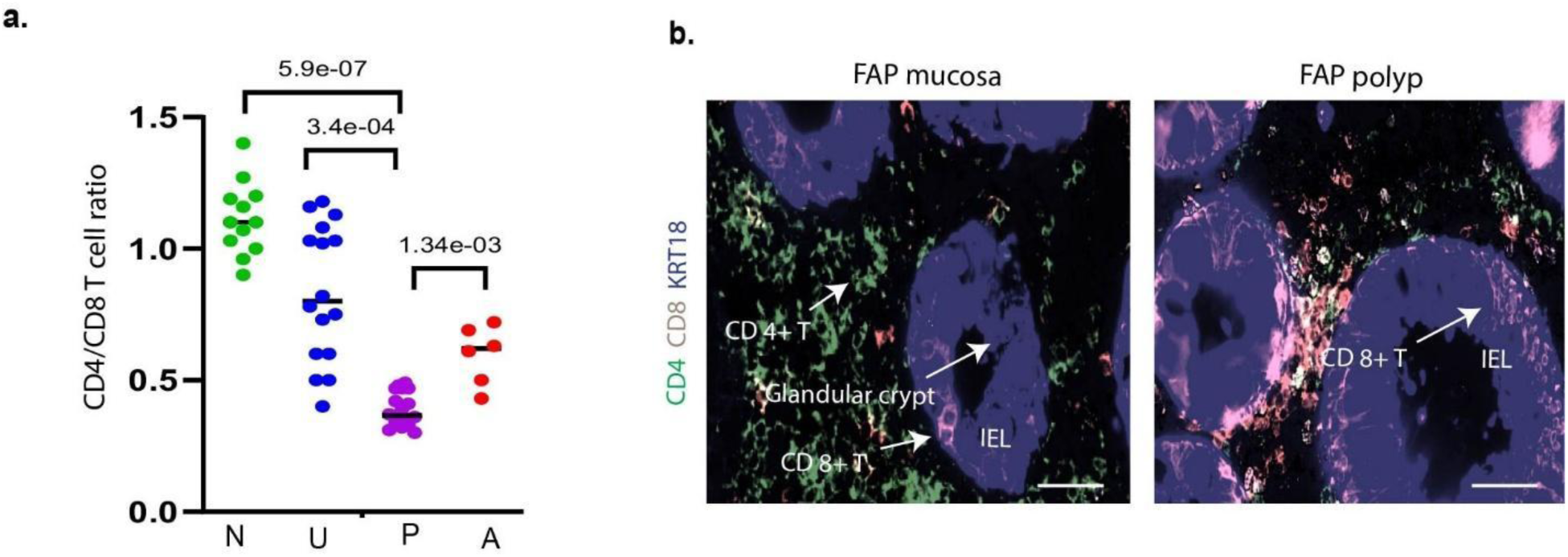
CD4+T and CD8+ T cell type composition changes across FAP disease continuum. (**a**) Scattered dot plots representing the CD4/BD8 T cell ratio across normal mucosa (N), FAP mucosa (U), FAP polyp (P) and, FAP adenocarcinoma/CRC (A). A two-tailed statistical t test was performed and adjusted P value has been indicated, (**b**) CODEX imaging showing CD8+ T cell tend to localize near the epithelial glandular crypts as intraepithelial lymphocytes (IEL). This is mostly observed in FAP polyps compared to FAP mucosa. Scale bar: 10µm

**Table S1:** Donor metadata. Twelve FAP and eight non-FAP donors consented for this study. See text for details.

| Donor study ID | Disease status | Current/ Age at collection | Sex | Race/Ethnicity | Cancer Status | Sample collected | Underlying health condition |
| --- | --- | --- | --- | --- | --- | --- | --- |
| A001 | FAP (clinical dx) | 50 / 45 | Male | Hispanic/Latino | Sigmoid colon cancer | Colectomy | Asthma |
| A002 | FAP (APC+) | 24 / 20 | Female | White | No cancer | Colectomy | Unknown |
| A014 | FAP (clinical dx, negative genetic test) | 25 / 22 | Female | white | No cancer | Colectomy | Childhood asthma |
| A015 | FAP (APC+) | 38 / 36 | Female | Hispanic/Latino | No cancer | Colectomy | Unknown |
| A018 | FAP (APC+) | 56 / 53 | Male | Hispanic/Latino | No cancer | Colonoscopy | Hypertension |
| A035 | FAP (clinical dx, negative genetic test) | 61 / 58 | Male | White | Desmoid Tumor | Colonoscopy | Hypertension |
| A040 | FAP | 14 / 12 | Male | Hispanic/Latino | No cancer | Colectomy | Unknown |
| A051 | FAP | 26 / 25 | Male | Hispanic/Latino | No cancer | Colectomy | Diabetes mellitus |
| A055 | FAP | 39 / 35 | Female | White | No cancer | Colectomy | Asthma |
| A057 | FAP (clinical dx, negative genetic test) | 54 / 52 | Male | Hispanic/Latino | No cancer | Colectomy | Unknown |
| A069 | Adenocarcinoma | 65 / 67 | Male | White | Colorectal cancer | Colectomy | Unknown |
| G | FAP | 35 / 32 | Male | Hispanic/Latino | Colorectal cancer | Colectomy | Unknown |
| CRC_15564 | Adenocarcinoma, medullary type | Unknown / 66 | Female | White | Colorectal cancer | Tumor | Unknown |
| CRC_8396 | Adenocarcinoma, medullary type | 34 / 31 | Male | Hispanic/Latino | Colorectal cancer | Tumor | Unknown |
| CRC_23135 | Adenocarcinoma, medullary type | Unknown / 45 | Female | White | Colorectal cancer | Tumor | Unknown |
| CRC_19779 | Adenocarcinoma, medullary type | 57 / 54 | Male | Hispanic/Latino | Colorectal cancer | Tumor | Unknown |
| B001 | Normal mucosa | 67 | Female | White | Healthy colon | Autopsy | Diabetes, hypertension |
| B004 | Normal mucosa | 78 | Male | Black | Healthy colon | Autopsy | Diabetes, hypertension |
| B005 | Normal mucosa | 24 | Female | White | Healthy colon | Autopsy | None |
| B006 | Normal mucosa | 38 | Male | White | Healthy colon | Autopsy | None |

**Table S2:**
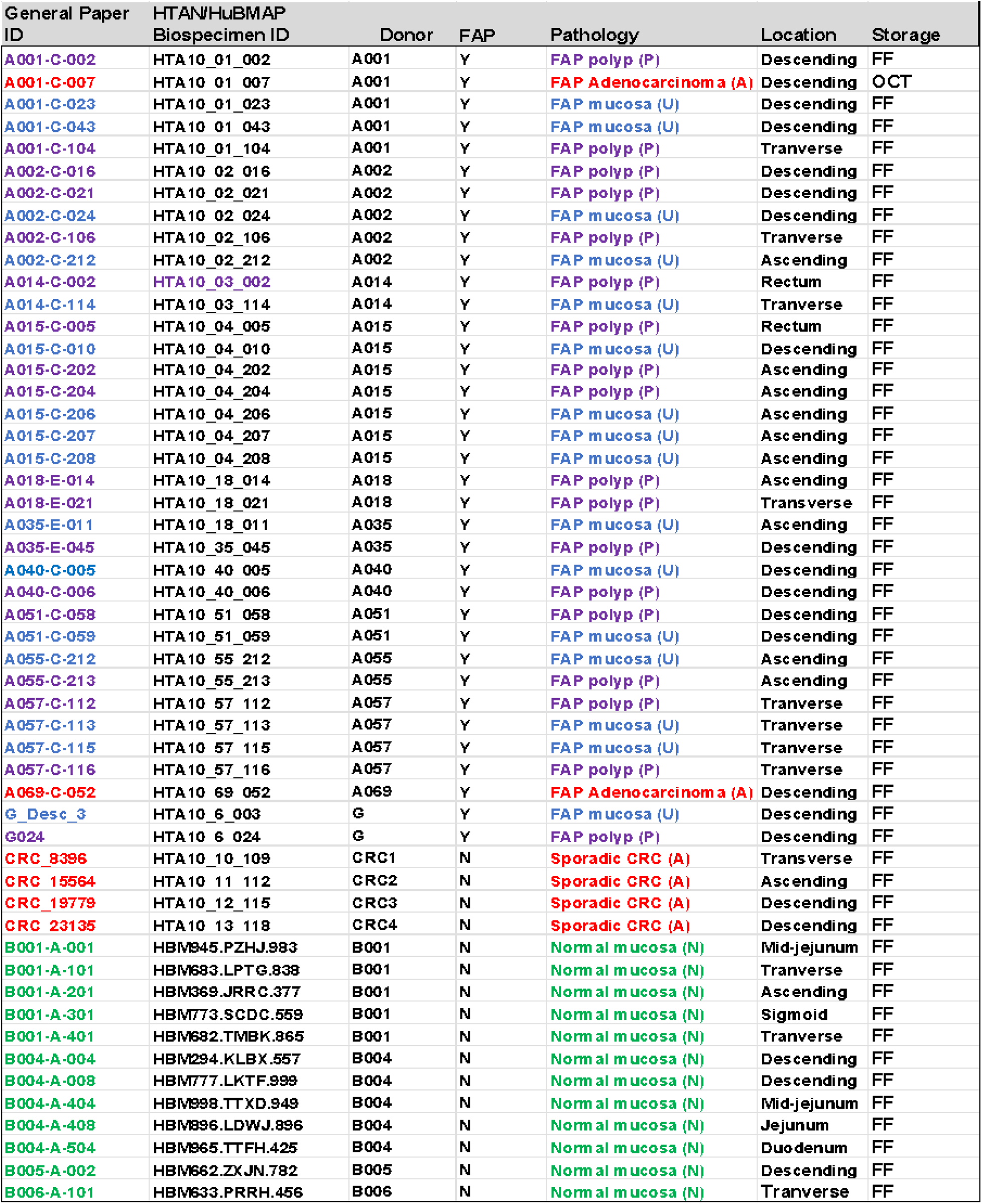
Sample metadata used in this study. Fifty-two samples including 12 normal mucosa, 16 FAP mucosa, 18 FAP polyp, 2 FAP adenocarcinoma and, 4 CRCs were selected for CODEX imaging and analyses. These samples are color coded (see text for details).

**Table S3.**
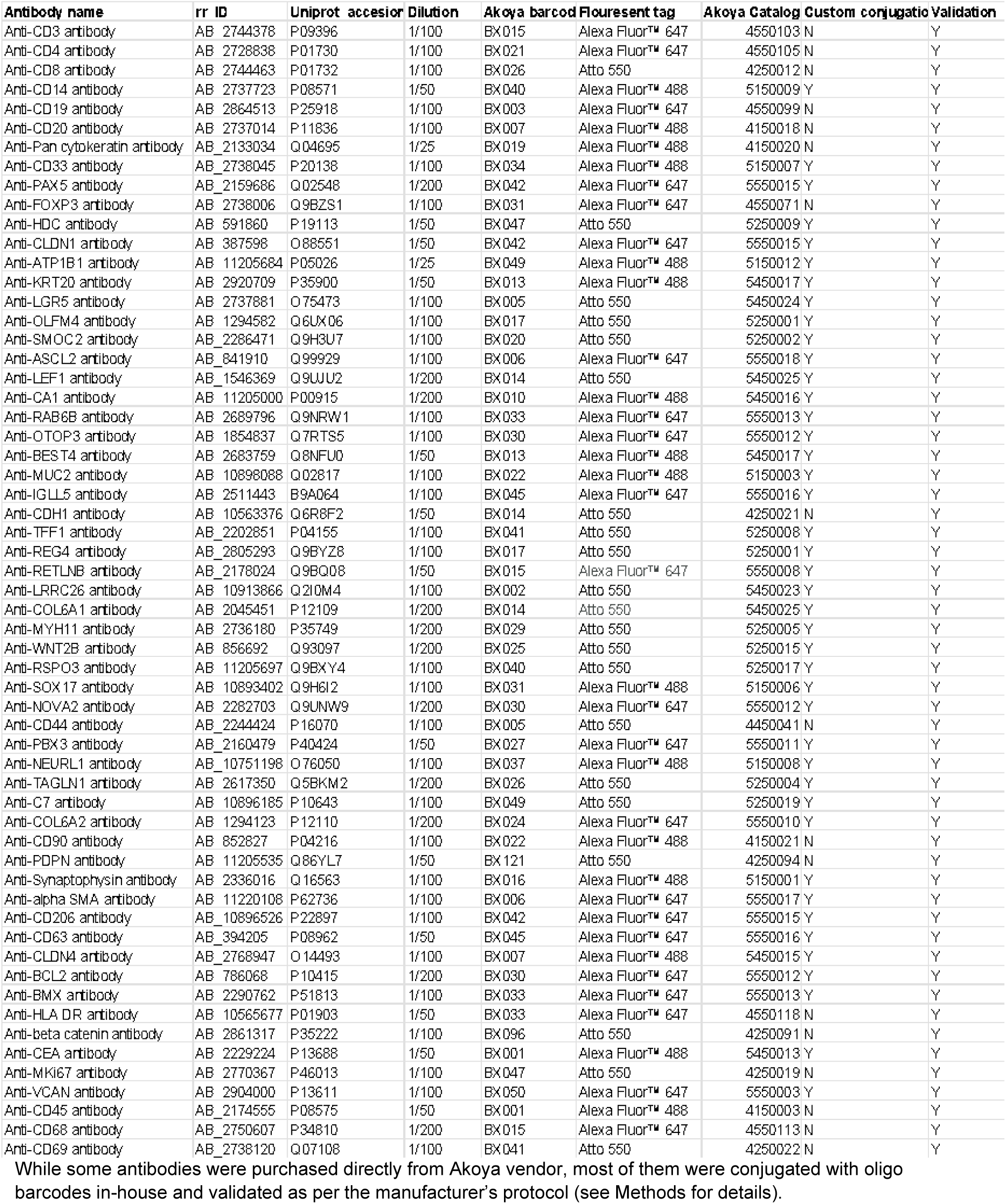
Details of CODEX oligo-barcoded antibodies used in this study. While some antibodies were purchased directly from Akoya vendor, most of them were conjugated with oligo barcodes in-house and validated as per the manufacturer’s protocol (see Methods for details).

